# Cities as evolutionary incubators for the global spread of the Spotted Lanternfly

**DOI:** 10.1101/2025.06.30.662460

**Authors:** Fang Meng, Anthony A. Snead, Aria Yang Zhang, Jason Munshi-South, Kristin M. Winchell

## Abstract

Habitat destruction and invasive species pose two of the greatest global threats to biodiversity. These factors do not operate in isolation, and nowhere is their interaction more apparent than in urban environments. Urban organisms rapidly evolve under novel ecological circumstances where they also encounter anthropogenic opportunities for range expansion. Here, we examine the role of urbanization in the invasive success of an emerging global pest, the Spotted Lanternfly, during colonization and expansion. We demonstrate that the invasive population in the United States has undergone three sequential bottlenecks, resulting in significantly reduced genetic diversity and elevated inbreeding. The success of this invasive population may be in part attributable to adaptation in the native range prior to the invasion: we detect divergence between urban and rural lanternflies in Shanghai, China, (the invasion origin) in genes related to stress response, metabolism, and detoxification pathways. Additionally, we detect genomic signatures of selection in the invasive population suggesting adaptive refinement as the invasion progresses. This study provides evidence of adaptive evolution in response to urbanization despite substantial loss of genetic diversity, and implicates adaptive responses to pesticide application, dietary shifts, and climate in the invasive success of the Spotted Lanternfly.

## Introduction

Biological invasions and habitat destruction pose two of the most significant threats to global biodiversity (Mack et al., 2000; Bellard et al., 2022). These processes are often studied in isolation, yet they interact in complex ways (Didham et al. 2007). Humans are rapidly transforming natural landscapes, creating globally distributed novel ecosystems typified by impervious surfaces and human infrastructure. In cities, species face novel selective pressures through urban heat island effects, human resource subsidies, and unique ecological niches (Szulkin et al., 2020) as well as encounter anthropogenic dispersal opportunities (Cadotte et al., 2017). Indeed, cities serve as primary hubs for biological invasions (Cadotte et al., 2017) and many non-native animals thrive in urbanized landscapes (Carlon & Dominioni 2024). Their success may be related to the global similarity of cities, enabling adaptive responses to urbanization in the native range to facilitate invasion despite genetic constraints in the founding population (Hufbauer et al., 2012, Borden & Flory, 2021). Evolution in urban environments may play a critical role in overcoming the challenges of introduction, and an underappreciated role in invasive spread.

The Spotted Lanternfly (*Lycorma delicatula*) provides an excellent opportunity to explore the interaction of urbanization and biological invasion. Native to China and South Asia, the Spotted Lanternfly was introduced to the United States in 2014 and has expanded from Pennsylvania throughout the northeast megalopolis. Research suggests the invasion originated from a single introduction from South Korea (Du et al., 2020; Kim et al., 2021). Mitochondrial DNA and microsatellite evidence identified Shanghai, China, as the probable source for the Korean invasion (Kim et al. 2021), and support a sequential invasion pathway from Shanghai to South Korea followed by a single introduction from South Korea to the United States (Du et al., 2020; Kim et al., 2021). Ten years later, Spotted Lanternflies now form massive swarms in cities from Philadelphia to New York City and beyond. Their expansion has been facilitated by human-mediated transport, absence of natural enemies, and the presence of the invasive tree-of-heaven (*Ailanthus altissima*), their preferred host plant (Ladin et al., 2023; Du et al., 2020). However, the role of adaptive evolution in their invasion success—in particular the role of urban-associated adaptive evolution—is not currently understood. We address this knowledge gap to better understand how adaptive evolution in cities facilitates biological invasions at both the colonization and expansion phases.

We analyze whole-genome sequences of Spotted Lanternflies from native (Shanghai, China) and invasive (northeastern United States) ranges to address three questions. (1) Does the invasive population exhibit reduced genetic diversity as expected from a single introduction? (2) Do historic and contemporary demography explain patterns of genetic diversity? (3) Has natural selection in the native and invasive range facilitated invasive success? Answers to these questions will advance our understanding of how invasive species overcome genetic constraints to rapidly adapt to novel environments, with implications for predicting and managing biological invasions in an increasingly urbanized world.

## Results

### Invasive US population exhibits reduced genetic diversity and elevated inbreeding

Given the proposed single introduction pathway supported by prior research (Du et al., 2020; Kim et al., 2021), we expected to detect genetic signatures of the founding event in the invasive population. Whole-genome sequencing of 118 *L. delicatula* from the northeastern USA (n=98) and Shanghai, China (n=20), reveals a substantial reduction in genome-wide heterozygosity in the invasive (USA) versus native (China) populations (Welch’s t-test: t=8.99, df=43.76, p-value=1.703e-11; Fig. 1a). Although genetic diversity is overall higher in China compared to the USA, the variance across individuals is greater in the USA (F-test: F=0.36, p=0.014). The observed difference in heterozygosity is robust despite unequal sample sizes and variance across locations (confirmed by bootstrap random subsampling: differences persist in 100% of iterations).

**Fig. 1.**
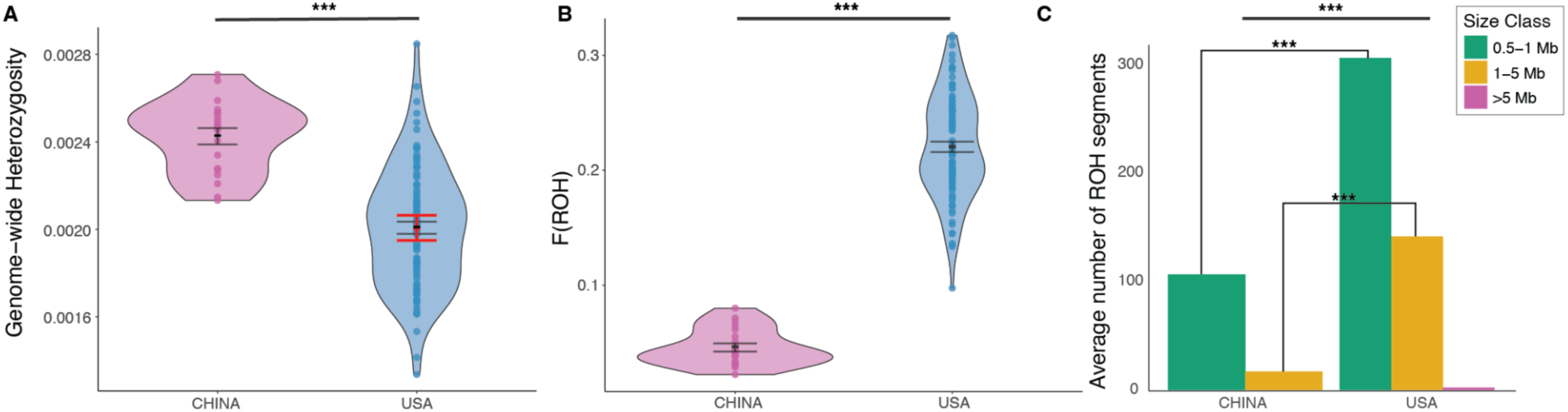
Genetic Diversity in the Invasive versus Native Ranges. **a,** Genome-wide heterozygosity in native China (n=20) compared to invasive USA (n=98) Spotted Lanternfly populations. Each dot represents an individual sample, with black lines indicating the mean and standard error of each population. The red point and lines indicate the mean and standard error of heterozygosity in 1,000 random subsamples of 20 USA individuals. **b,** Violin plots showing the distribution of F_ROH_ as a measure of inbreeding in China and USA populations (p < 2.2e-16). Each dot represents an individual sample, with black lines indicating the mean and standard error. **c,** Barplots displaying the average number of runs of homozygosity (ROH) segments per individual for each size class (0.5–1 Mb, 1–5 Mb, >5 Mb) in China and USA populations. Significance levels: p<0.001 ***.

We detected signatures of inbreeding in the invasive population via runs of homozygosity (ROH). Overall, USA individuals exhibited significantly higher inbreeding coefficients (FROH) compared to Chinese individuals (Welch’s t-test: t=-30.22, df=85.04, p<2.2e-16; Fig. 1b). USA individuals harbored nearly four times as many ROH segments (Fig. 1c; Welch’s t-test: t=-28.20, df=48.88, p<2.2e-16). Small ROH segments (0.5–1 Mb) were overrepresented in the USA compared to China (Welch’s t-test: t=-22.54, df=35.76, p<2.2e-16) and medium ROH segments (1–5 Mb) were eight times more abundant in the USA (Welch’s t-test: t=-28.38, df=115.38, p<2.2e-16). Notably, we identified 83 long ROHs (>5 Mb) in 44 individuals in the USA, but long ROHs were entirely absent in the native-range (China) samples. While the high frequency of small ROHs (0.5–1 Mb) could reflect ancient inbreeding (Kirin et al., 2010), their overrepresentation in the recently established USA population more likely reflects identity-by-descent from a founding event (Stoffel et al., 2021). Moreover, the overrepresentation of medium ROHs and presence of especially long ROHs in the USA suggest recent shared ancestry and intense inbreeding among individuals (Kirin et al., 2010; Ceballos et al., 2018; Kardos et al., 2017), consistent with founder effects during invasion (Uller & Leimu., 2008).

Long RoHs can arise by both demographic processes and natural selection. Given the absence of long ROHs in the native range, extended homozygous regions in the USA more likely result from demographic processes such as recent bottlenecks or founder effects. Similar patterns of elevated FROH were documented in the Common Wall Lizard (*Podarcis muralis*), where introduced populations showed substantially higher proportions of long ROHs (>1MB) compared to the native source population (Bode, 2025). While we cannot rule out selection, the overall pattern we observe is more consistent with demographic history, with limited evidence of selection at locations within long ROH (Fig. S2). Future work incorporating recombination maps or haplotype-based analyses would help disentangle these effects and clarify whether long ROHs reflect adaptation or stochastic demographic events.

### Historic and contemporary demography explain reduced genetic diversity in the invasive range

To understand patterns of reduced genetic diversity in light of founder effects and bottlenecks, we reconstructed demographic history and assessed current population differentiation. We observed higher linkage disequilibrium (LD) across all physical distances in the USA (r² ≈ 0.15 at 300 kb vs. r² ≈ 0.08 in China; Fig. 2a), consistent with a recent founder effect (Reich et al., 2001; Pritchard & Przeworski, 2001). As expected, LD decayed more rapidly in native populations, characteristic of larger and more stable effective population sizes (Slatkin, 2008; Flanagan et al., 2021). Nonlinear regression yielded identical recombination rates (1 cM/Mb) for both populations, suggesting LD differences are driven by demography. Best-fit model parameters (China: d₀ = 0.8, b = 0.1; USA: d₀ = 0.8, b = 0.2) show elevated background LD in the USA, supporting a recent bottleneck.

**Fig. 2.**
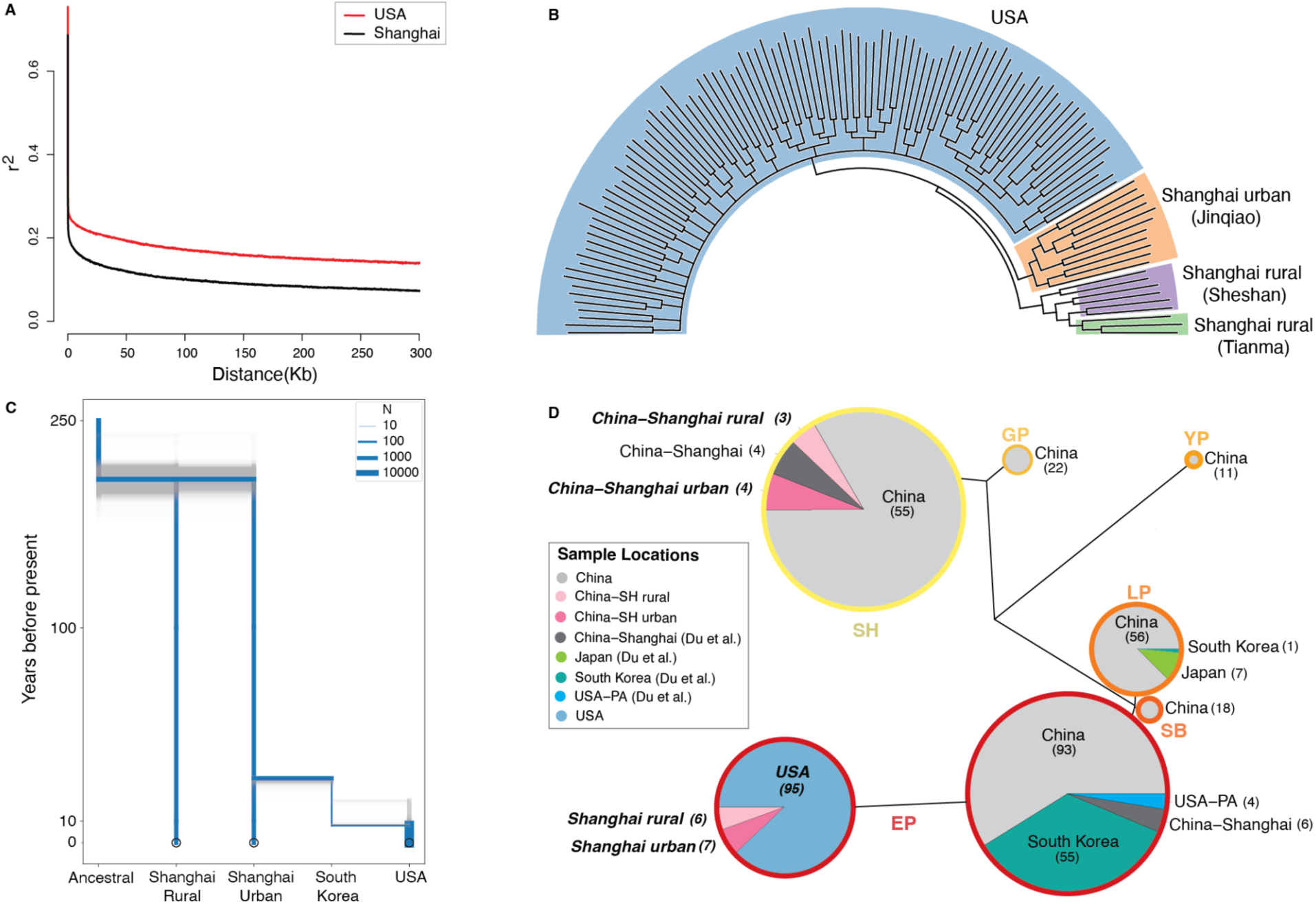
Demographic History of the Spotted Lanternfly Invasion. **a,** Decay of linkage disequilibrium (LD) with maximum distance of 300 kb between SNP pairs in Shanghai (black) and USA (red) populations. **b,** Maximum-likelihood phylogenetic tree based on 158,123 SNPs showing evolutionary relationships among 118 Spotted Lanternfly individuals. Blue branches represent USA populations, orange represent Shanghai urban, and green and purple represent the two Shanghai rural populations. Individuals from USA and urban Shanghai share a recent common ancestor, and this clade is sister to the individuals from the Shanghai rural locations. **c,** Topologies for the best demographic model (Independent Urban Bridgehead): Common Ancestral → Shanghai rural/urban (parallel), Shanghai urban → Intermediate Population (potentially South Korea)→ USA. Time is in years, with a generation time of 1 year. The width of the branches indicates effective population sizes. Shaded regions represent bootstrap replicates from demographic model fitting and illustrate uncertainty in parameter estimates. **d,** Mitochondrial haplotypes of 510 Spotted Lanternfly individuals across the native and invasive ranges, combining samples from the present study with those from Du et al. (2021) (samples from the present study are indicated by bold and italic text). Haplotype groups correspond to those named by Du et al. (2021), colored by the outside circle (note EP is split into two clusters but both are nested within the EP haplotype group; see Fig. S3 for full mtDNA tree with haplotype assignment). Circle size corresponds to number of individuals per cluster, with sample sizes per location provided.

Contemporary population structure also reflects this bottleneck. Phylogenetic inference recovered three monophyletic clades corresponding to the USA and two Chinese populations (Shanghai urban and rural; Fig. 2b). The invasive USA population shares more recent ancestry with urban than rural Shanghai, consistent with an urban origin (though the precise source could lie in an unsampled location within that range). Mitochondrial DNA analysis confirms that USA samples cluster within the East Plain (EP) lineage identified by Du et al. (2020) as the invasion origin. Both EP and Southeast Hills (SH) haplotypes are present in urban and rural Shanghai, indicating mtDNA is not segregated by environment (Fig. 3d, Fig. S3).

**Fig. 3.**
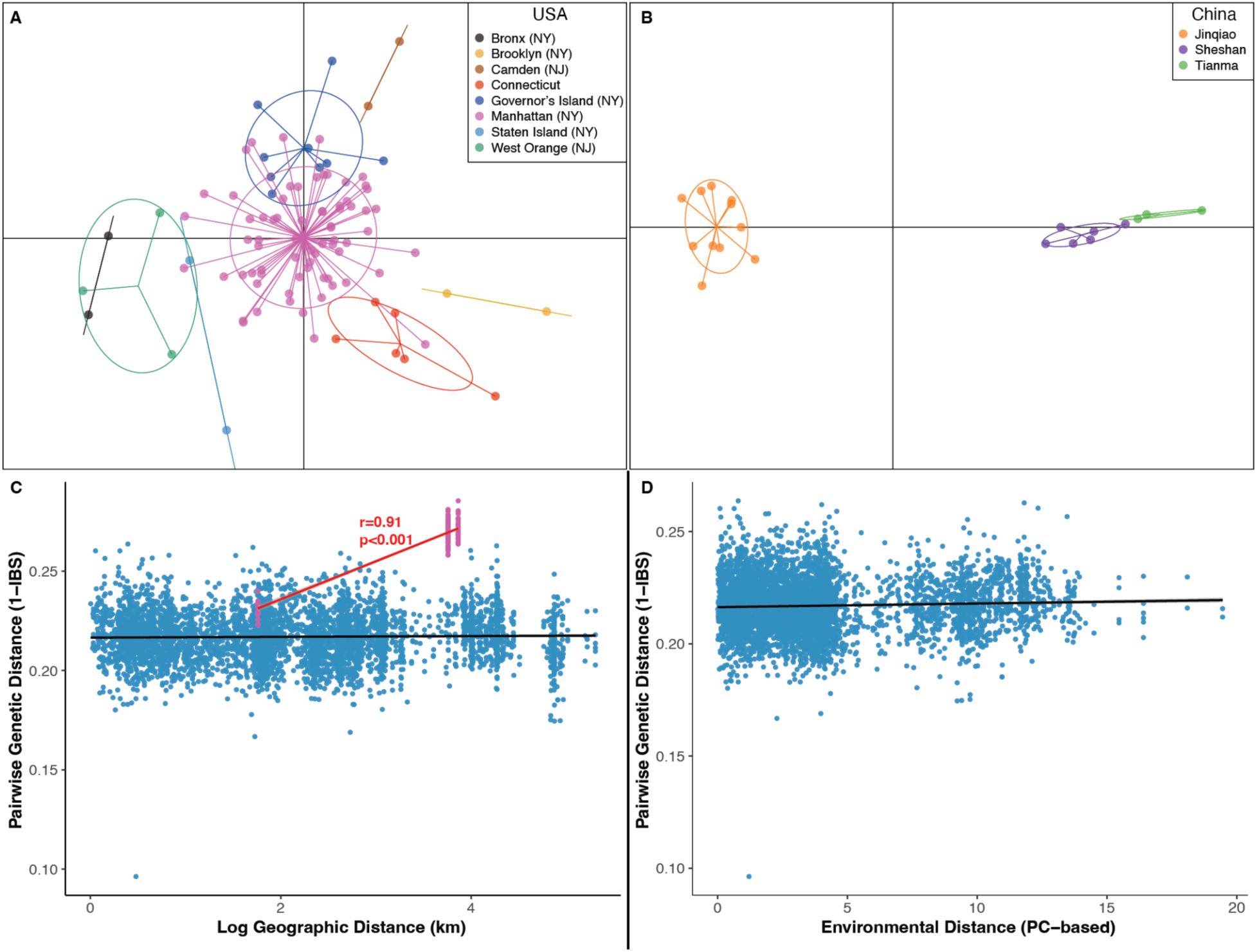
Population structure and environmental associations in native and invasive Spotted Lanternfly populations. **a,** Discriminant Analysis of Principal Components (DAPC) plot for 98 USA individuals spanning 0– 200km geographic distance. Each point represents an individual, colored by sampling location, with ellipses indicating 95% confidence intervals around population means. **b,** DAPC plot for 20 individuals from Shanghai urban (Jinqiao) and rural (Sheshan and Tianma) locations spanning 0–40km distance. **c,** Isolation-by-distance analysis showing contrasting patterns between native and invasive populations: Shanghai populations (pink points, red line) exhibit strong correlation between pairwise genetic distance (1-IBS) and geographic distance (log-transformed, km), while USA populations (blue points) show no significant relationship. **d,** Isolation-by-environment analysis showing no significant relationship between genetic differentiation and environmental distance for USA populations. Lines in C and D represent fitted linear regressions.

We next used momi2 to infer historical population size changes and divergence. Our best-fitting model (AIC weight = 0.936; Fig. 2c, Tables S2–S3) supports a sequential invasion involving three population bottlenecks. The first, 171.4 ± 7.0 years ago (range: 149.9–226.0), marks divergence between urban and rural Shanghai, coinciding with Shanghai’s rapid urbanization. The second, 29.7 ± 0.86 years ago (range: 19.6–30.0), aligns with the species’ detection in South Korea in 2004 (Kim et al., 2021). The third, most recent, at 8.95 ± 2.97 years ago (range: 8.0–20.0), coincides with its detection in Pennsylvania in 2014 (Barringer et al., 2015). This timing is also supported by ROH-based coalescent analysis, where long ROHs in the USA suggest common ancestry ∼8.9 generations ago (Table S4), assuming a recombination rate of 1 cM/Mb (Foote et al., 2021).

These results support a sequential “stepping-stone” invasion model (Estoup & Guillemaud, 2010), in which the urban Shanghai population gave rise to a bridgehead population (presumably South Korea; Kim et al., 2021; Du et al., 2020) that subsequently founded the USA population. While sampling from additional source populations is needed to confirm this pathway, the strong alignment between inferred bottlenecks, historical records, phylogenetic relationships, and ROH-based estimates provides compelling support. Together, these demographic events explain the markedly reduced genetic diversity observed in the invasive USA population.

Despite samples in the USA spanning a considerable geographic distance (0–200km), we did not detect significant population differentiation (Fig. 3a), with all individuals forming a single cluster of shared ancestry as determined by both ADMIXTURE and sNMF (K=1, Fig. S5). We detected no correlation between genetic and geographic (Mantel test: r=0.011, p=0.419; Fig. 3c), or environmental distances (Mantel test: r=0.075, p=0.075; Fig. 3d), demonstrating neither spatial proximity nor environmental similarity strongly influences population structure in the invasive range. In contrast, we observe genetic structure across urban and rural locations in Shanghai despite being separated by only 30–40 km (Fig. 3b), with a single ancestral population still best supported by ADMIXTURE and sNMF (K=1, Fig. S5) suggesting recent population differentiation rather than a deep ancestral split between populations. Shanghai populations show strong isolation-by-distance (Mantel test: r=0.91, p<0.001; Fig. 3c), although we cannot disentangle the effects of isolation-by-distance versus environment with our sampling structure. These results support limited gene flow over short distances in the native range, consistent with the localized movement patterns and limited flight capability reported for Spotted Lanternfly (Wolfin et al., 2019; Kim et al., 2021).

The weak genetic structure in the USA is notable given the strong differentiation observed in Shanghai across smaller spatial scales, suggesting that limited population differentiation is not an intrinsic biological characteristic of the species. The lack of population differentiation in the USA likely stems from four complementary factors. First, the invasive population may not have had sufficient time to reach migration-drift equilibrium since introduction (Wang & Bradburd, 2014). Second, founder effects may have reduced genetic diversity to the point that little variation remains to differentiate populations. Third, human-facilitated dispersal may enable long-distance population connectivity through transportation networks (Ladin et al. 2023), although human-mediated transport is unlikely to be the primary reason for the population structure differences as Shanghai also has an extensive transportation infrastructure. Fourth, ecological pressures from natural enemies and competitors could constrain population growth and dispersal in the native range but not in the invasive range where these pressures are absent.

### Natural selection may be facilitating the Spotted Lanternfly invasion

An outstanding question in invasion biology is how do introduced species overcome genetic constraints imposed by founder events to thrive in novel environments (Estoup et al., 2016; Sax et al., 2007)? Population bottlenecks should limit adaptive evolution by reducing genetic variation and increasing the effect of drift in small founding populations (Barrett & Schluter, 2008). Many species overcome the challenges of colonization through multiple introductions and gene flow among subpopulations resulting in similar (or even higher) levels of diversity in the invasive range (Kołodziejczyk et al. 2025), but this is not the case with the Spotted Lanternfly in the USA. The demographic history of the Spotted Lanternfly invasion, with multiple bottlenecks and reduced genetic diversity, could limit adaptive responses as the invasion progresses. Nevertheless, the Spotted Lanternfly is abundant in the USA, particularly in cities. One explanation for this apparent paradox is that selection in the native range may have favored phenotypes that are adaptive in urban environments generally (i.e., the Anthropogenically Induced Adaptation to Invade hypothesis; Hufbauer et al. 2012, Borden and Flory 2021). Upon introduction to the northeast megalopolis of the USA, these individuals would already be adapted to urban living. Natural selection may have then acted to refine adaptive responses to the subtly different selection landscape in the USA. To evaluate this hypothesis, we examined genomic signatures of natural selection in the native and invasive ranges.

Across three locations in Shanghai (one urban, two rural), we identified 760 SNPs under selection using a PCA-based genome scan. Among the most significant outliers were 21 functionally annotated genes, many with roles in immune response, metabolism, stress tolerance, and developmental processes (Fig. 4a). We conducted Population Branch Excess (PBE) analyses to identify genomic regions under selection in urban versus rural locations (Fig. 4b). Genes with urban-biased selection include genes are implicated in functions that are likely important in urban environments such as reproductive processes (RAB1A; Liu et al., 2024), pathogen defense (TASP1; Qu et al., 2018; Kaur & Roberts, 2024), and wing development (LEMD3; Wagner et al., 2010). Genes with rural-biased selection primarily reflect functions involved in cellular maintenance and regulation, including chromatin architecture (CHD5), energy metabolism (GYG1), cytoskeletal dynamics (FHOD1), and extracellular matrix formation (EXTL3). Several SNPs in these genes have contrasting patterns of allele frequency between urban and rural populations (Fig. 4c), providing insight into directional selection in urban Shanghai and suggesting possible targets of natural selection that facilitated invasion success.

**Fig. 4.**
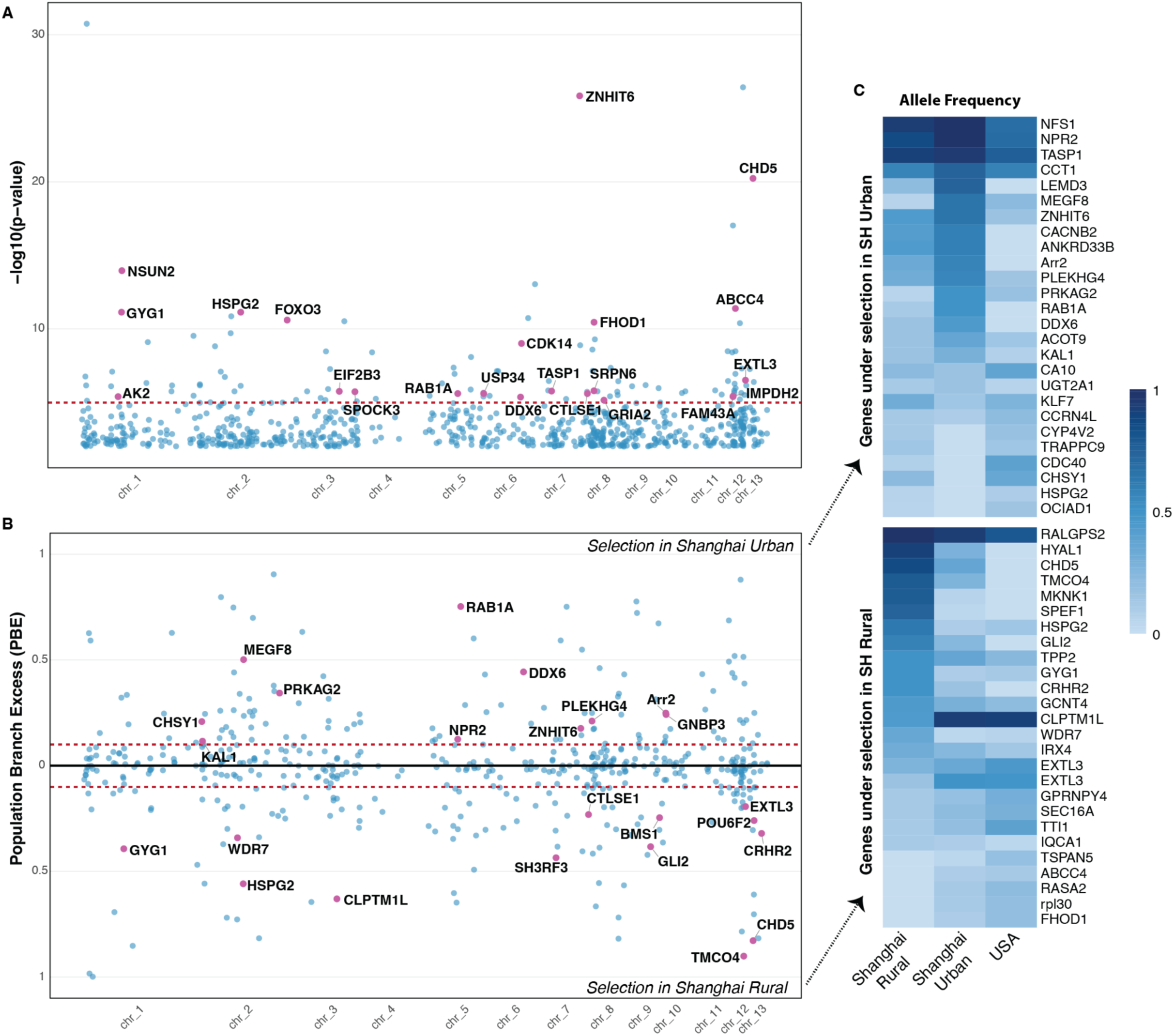
Signatures of selection in the native range Genome-wide patterns of differentiation between urban and rural locations in Shanghai, China. **a,** Manhattan plot showing outliers detected using PCA-based genome scans. The dashed red line indicates the significance threshold (α<1×10–5 with FDR-corrected p<9.7e-06) for top candidate SNPs. Outliers above this threshold with functional gene names are represented by pink points, while blue points indicate all remaining 760 candidate SNPs (FDR<0.01). **b,** Mirrored Manhattan plot of Population Branch Excess (PBE) differentiating outliers specific to Shanghai urban (positive values, upper panel) and rural (positive values, lower panel) populations. Dashed red lines mark thresholds (±0.1 PBE). Pink highlighted points indicate significant SNPs annotated with functional gene names. **c,** Heatmaps of allele frequency for outlier SNPs in annotated genes identified by PBE in urban (top) or rural (bottom) populations in Shanghai, China. Colors represent allele frequencies ranging from low (light blue) to high (dark blue) across Shanghai rural, Shanghai urban, and invasive USA populations.

We also detected several genes exhibiting selection signatures in both urban and rural habitats, including ABCC4, CTLSE1, NSUN2, and HSPG2. Of these, ABCC4 plays a critical role in insecticide resistance and detoxification of plant secondary metabolites in arthropods (Dermauw & Van Leeuwen, 2014; Wu et al., 2019). This finding is particularly relevant given targeted insecticide application by municipal authorities in China in areas with high Spotted Lanternfly densities, especially city parks (Shanghai Botanical Garden, 2018; Tianjin Municipal Bureau of City Management, 2016). Additionally, selection on ABCC transporters could be related to detoxification of toxic compounds in their preferred host plant, *Ailanthus altissima* (tree-of-heaven), as these insects sequester ailanthone and other quassinoids for defense (Song et al., 2018). Selection on detoxification genes across both urban and rural environments suggests that xenobiotic metabolism—related to insecticides, diet, or both—remains a key adaptive challenge regardless of habitat type.

In the USA population, we identified SNPs under selection using partial RDA (which explicitly incorporates environmental variables and spatial autocorrelation) and a PCA-based genome scan (which detects differentiation among individuals agnostic to environmental variation or spatial distribution). We identified 7,318 outlier SNPs using PCAdapt (Fig. 5a) and 2,675 outlier SNPs through pRDA analysis (Fig. 5b-c). In the pRDA, temperature was the primary association with outlier SNPs (1,262 SNPs, 47.2%), followed by precipitation (847 SNPs, 31.7%) and urbanization (566 SNPs, 21.2%). We identified a conservative set of 184 candidate SNPs as those detected as outliers by both methods (Fig. 5a-c). The chromosome-wide differentiation across chromosomes 1, 6, and 8 is not attributable to elevated LD or SNP-density bias on these chromosomes (Fig. S9, Table S5). Similar chromosome-wide differentiation has been reported using PCA-based methods (e.g., Bekele et al., 2018), attributable to selection on linked haplotypes or large-effect genomic features. Linked haplotypes are unlikely to explain this pattern in our data given the similar LD among chromosomes (Fig. S8), although we note that chromosome 1 has the greatest number and longest total lengths of long ROHs. Severe founder effects may lead to chromosomal regions represented by few ancestral copies, providing little genetic variation for recombination to act upon. Additionally, elevated inbreeding increases the probability of inheriting two copies that are identical by descent. Reduced recombinational breakdown of identity-by-descent blocks, rather than selection, could contribute to the chromosome-wide signals we observe, although is unlikely to explain the observed patterns fully.

**Fig. 5:**
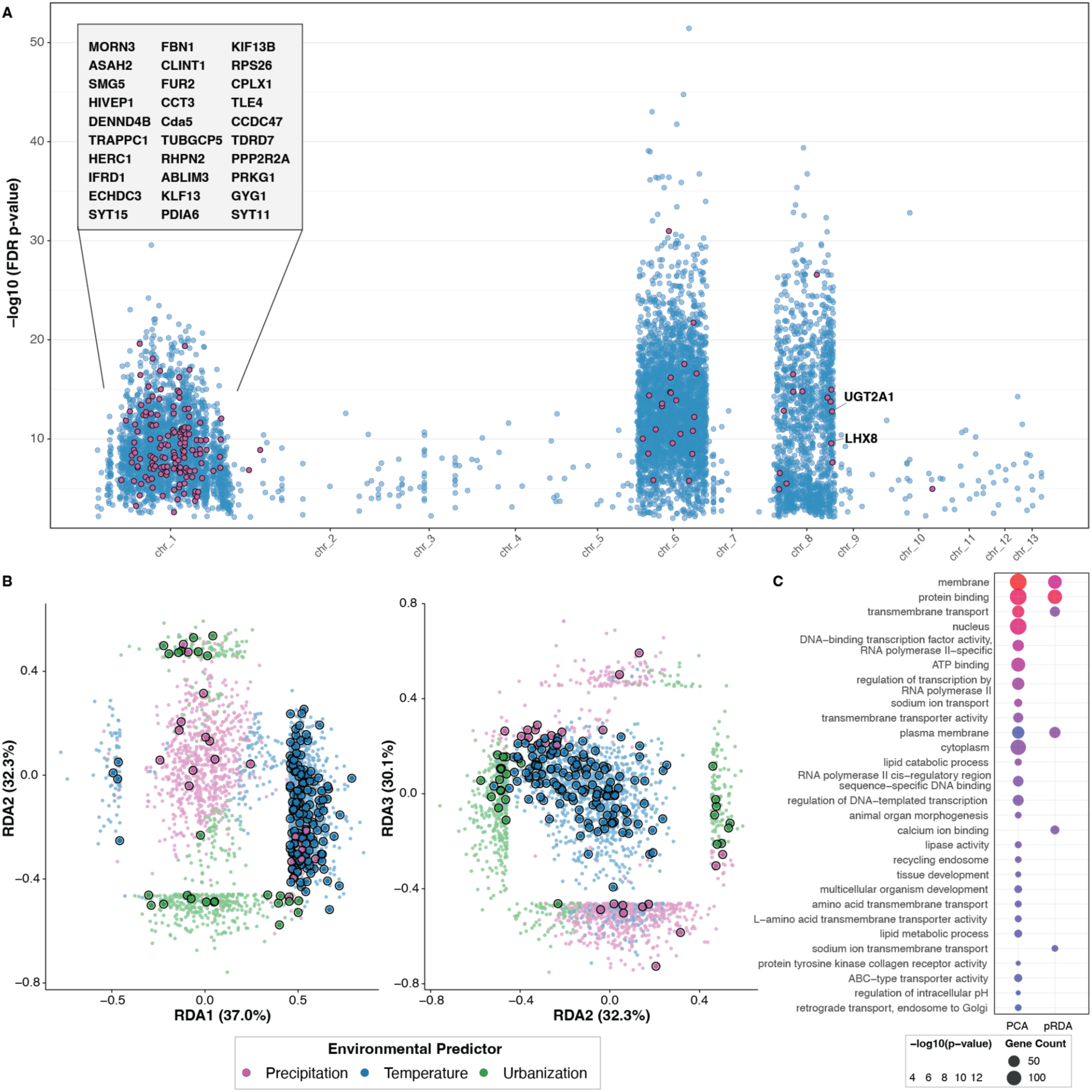
Signatures of selection in the invasive range. **a,** Manhattan plot displaying selection signals across chromosomes. Blue dots represent significant SNPs (FDR α < 0.01, n = 7349) detected by PCAdapt, with pink dots (n = 184) indicating SNPs identified by both PCA and RDA methods. FDR-adjusted p-values are from the K=6 PCAdapt run. Gene names are shown for key candidate genes, with those on chromosome 1 appearing in the box above CHR1. **b,** RDA ordination plot showing the distribution of selection candidates along the first (RDA1, 37.0%) and second (RDA2, 32.3%) environmental axes, and second (RDA2, 32.3%) and third (RDA3, 30.7%) environmental axes. SNPs are colored according to their primary environmental association (blue: temperature; purple: precipitation; green: urbanization; black highlight: SNPs detected by both methods). **c,** Bubble plot of significantly enriched Gene Ontology (GO) terms across different selection detection methods. The horizontal axis indicates method specificity (PCAdapt-only: 26 terms; pRDA-only: 6 terms; Overlap: 4 terms). Bubble size represents gene count within each GO category, and color intensity corresponds to statistical significance (–log10(p-value)).

Candidate SNPs identified by both methods are located in 32 functionally annotated genes. These include several genes involved in lipid and carbohydrate metabolism, including ECHDC3, GYG1, and ASAH2 (Yan et al., 2021; Zhang et al., 2017; Zhou et al., 2013). We also identified genes with potential roles in environmental stress response, such as PDIA6, which was associated with temperature in our pRDA and is upregulated under oxidative and thermal stress in honeybees (Meng et al., 2021). Notably, PRKG1, the ortholog of the *Drosophila* foraging gene, was associated with precipitation. This gene modulates foraging behavior and cGMP signaling (Ben-Shahar et al., 2002) and also directly controls cytochrome P450-mediated detoxification capacity, which may be relevant to xenobiotic metabolism (Amichot & Tarès, 2021). We identified several other genes involved in neural signaling and synaptic transmission, including CPLX1 (associated with precipitation) and SYT11/SYT15 (associated with temperature) (Takai et al., 2020; Wang et al., 2023). Adaptive evolution across stress response, metabolic, and neural signaling pathways may collectively contribute to the species’ rapid establishment and spread across the northeastern United States.

We conducted Gene Ontology (GO) enrichment analysis on candidate gene sets identified by PCA and pRDA approaches separately. Only four GO terms were significantly enriched in both gene sets: membrane, protein binding, transmembrane transport, and plasma membrane components. Candidate genes identified by only PCA were enriched for developmental and metabolic pathways such as organ morphogenesis, tissue development, multicellular organism development, lipid metabolism, and ATP binding (Fig. 5d). Functional enrichment in pRDA-specific candidates highlights adaptation to environmental variation in the invasive range with two significantly enriched terms: calcium ion binding and sodium ion transmembrane transport (Fig. 5d). Calcium ion binding plays crucial roles in insect cold sensing, and calcium signaling mediates rapid cold-hardening responses in insect tissues (Teets & Denlinger, 2013). Given the northward expansion of the invasion into colder climates, this functional enrichment may reflect adaptation to seasonal cold and thermal variability as the invasion progresses. Enrichment for sodium ion transmembrane transport is equally intriguing, as it maintains neural function across temperature gradients and potentially confers resistance to pesticides such as pyrethroids (Dong et al., 2014). Pesticides have been the main approach to Spotted Lanternfly control in the USA as the invasion has unfolded (Urban and Leach 2023), and the functional enrichment we find here may indicate an adaptive response to this selective pressure.

Lastly, we identified five genes under selection in both Shanghai and USA populations, highlighting parallel responses to urbanization in the native and invasive range. Of these, UGT2A1 (UDP Glucuronosyltransferase Family 2 Member A1) stands out as under urban-specific selection by our PBE analysis in Shanghai and also associated with urbanization in our pRDA in the USA. UGTs enhance the solubility and excretion of plant allelochemicals and synthetic compounds such as insecticides (Li et al., 2007; Ahn et al., 2012), pointing to adaptation to pesticide application or host-switching in urban environments in both the native and invasive range. Although the Spotted Lanternfly specializes on the tree of heaven, it also readily consumes a large variety of other plant species that it may rely on more substantially as it invades environments where tree of heaven is less abundant (Barringer and Ciafré 2020).

Despite exhibiting significant genome-wide reduction in genetic diversity, observed heterozygosity (Ho) at the 184 SNPs identified under selection in the invasive population remained comparable to that in the native populations (USA: 0.210±0.017, Shanghai rural: 0.211±0.017, Shanghai urban: 0.20±0.015) (Fig. S10). We verified this pattern was not an artifact of uneven sampling) finding no significant change in observed heterozygosity if sample sizes were comparable to those in China based on 100 replicate subsamples of 10 individuals (Kruskal-Wallis: χ²=2.50, df=2, p=0.286). This pattern aligns with theoretical predictions and empirical findings that quantitative variation relevant to fitness is often less affected by bottlenecks than neutral marker diversity (Dlugosch & Parker, 2008; Bock et al., 2015). The disconnect between molecular marker diversity and adaptive potential is expected to be particularly pronounced where traits are polygenic and under strong directional selection (Estoup et al. 2016). While we find a substantial reduction in genome-wide genetic diversity following the sequential invasion bottlenecks, diversity at loci under selection appears well-preserved. Natural selection can maintain functional variation at ecologically relevant loci even as genome-wide diversity declines during bottlenecks (McKay & Latta, 2002; Merilä & Crnokrak, 2001). The candidate genes we identified are associated with stress response, metabolism, and detoxification pathways critical for survival in novel environments, representing precisely the kind of variation expected to be under selection during invasion (Kingsolver et al., 2012). The preservation of functional variation at these loci demonstrates how the evolutionary potential of an invasive population can remain intact despite significant reductions in genome-wide diversity.

## Conclusion

Here we reveal a complex invasion history of the Spotted Lanternfly characterized by repeated bottlenecks yet ongoing selection despite significantly reduced genetic diversity. These findings challenge the traditional genetic paradox of invasion, demonstrating successful establishment and rapid spread despite apparent genetic constraints. Through our analyses, we find that adaptation to urban environments may hold the key to understanding this paradox in the Spotted Lanternfly. Through adaptation to urban environments in the native range, colonizing individuals may have already possessed adaptations relevant to success in South Korea and the eastern United States, both of which are highly urbanized at the epicenter of the invasions. Urban-specific selection signatures in the native range were detected in genes related to stress tolerance, xenobiotic metabolism, and energy regulation. This finding raises the intriguing possibility that adaptation to pesticide application and host-plant variation in the native range primed the species to tolerate other urban environments, as predicted by the Anthropogenically Induced Adaptation to Invade hypothesis (Hufbauer et al. 2012, Borden & Flory 2021). Ecological filtering on these exaptations may have been subsequently followed by adaptive refinement in the invasive range as the lanternflies were exposed to novel ecological pressures. We detect signatures of ongoing selection in the invasive range in detoxification genes as well as genes related to thermal and hydric stress, suggesting that limited genetic variation can still produce adaptive responses in novel environments. As global connectivity intensifies along with human population growth, the synergistic effects of urbanization and biological invasion are increasingly important to understand. Evolutionary adaptation likely plays a critical role in the success of invasive species, and one that must not be overlooked if we aim to predict and manage biological invasions in an increasingly urban world. Only by studying invasion and urbanization as interconnected parts of a whole can we really interpret and understand the resulting eco-evolutionary patterns that emerge at their interface. This holistic perspective may be the key to understanding the success of the Spotted Lanternfly and other rapidly spreading global pests. Indeed, cities may act as evolutionary incubators, setting the stage for the global spread of invasive species.

## Methods

### Sample Collection and Sequencing

We collected adult Spotted Lanternflies from both invasive and native ranges to investigate genomic patterns associated with the USA invasion. In the invasive range, specimens were collected opportunistically between October 6th and November 7th, 2022, across Pennsylvania, New Jersey, New York, and Connecticut, with intensive sampling in New York City. Manhattan was strategically targeted due to its highly urbanized landscape and convergence of multiple transportation networks, potentially facilitating colonization from diverse source populations. We selected 98 specimens for whole-genome sequencing across the sampled area. To establish baseline genetic metrics from the native range, we collected 20 additional adult Spotted Lanternflies in September 2023 from Shanghai, China. This native range sampling incorporated both urban and rural environments, with 11 individuals from Shanghai’s city center and 9 from two rural locations (Fig. S1).

For USA samples, we extracted genomic DNA from thoracic and head tissues using the Qiagen DNeasy Blood & Tissue Kit following manufacturer protocols. DNA quantification was performed using the Qubit 1X dsDNA High Sensitivity Assay. Library preparation employed the NEBNext® Ultra™ II FS DNA Library Prep Kit with unique barcoding via NEBNext® Multiplex Oligos. We validated library quality and concentration using the 4200 TapeStation system. Sequencing was performed on the Illumina NovaSeq 6000 platform with 150 bp paired-end reads, achieving a minimum coverage of 5× per sample. For Shanghai samples, we extracted genomic DNA using either the PureLink® Genomic DNA Mini Kit or the UE Genomic DNA Small Volume Kit. Library preparation and sequencing were conducted by JMDNA Bio-Medical Technology Co., Ltd. (Shanghai) on the Illumina NovaSeq X Plus platform, generating a minimum coverage of 3× coverage per sample.

### Sequence Processing

Raw reads were basecalled using *Picard IlluminaBasecallsToFastq* version 2.23.8 (Broad Institute, 2019), with APPLY_EAMSS_FILTER set to false. Following basecalling, the reads were demultiplexed using *Pheniqs* version 2.1.0 (Galanti, Shasha, & Gunsalus, 2021). The entire process was executed using a custom Nextflow pipeline, *GENEFLOW* (New York University Genomics Core, 2023). We then processed sequencing data using the Variant Calling Pipeline with GATK4 (gatk/4.3.0.0), which follows the Broad Institute’s best practices for variant discovery (DePristo et al., 2011). Initial quality control was performed using FastQC v0.11.9 (Andrews, 2010). Raw reads were trimmed for adapters and low-quality bases with Trimmomatic v0.39 (Bolger et al., 2014) and aligned to the reference genome that masked the repetitive region (Snead et al., 2025) using BWA v0.7.17 (Li & Durbin, 2009). Post-alignment, duplicate reads were marked with Picard’s MarkDuplicates (picard/2.17.11), and reads were sorted using samtools v1.14 (Danecek et al., 2021.a). For variant discovery, we employed GATK HaplotypeCaller (v4.3.0.0) to generate gVCF files for each individual, followed by joint genotyping across all 118 samples from both native and invasive ranges using GenotypeGVCFs. This joint calling approach enabled a comparison of genetic variation between Shanghai and U.S. populations while maximizing our ability to detect shared and population-specific variants. We obtained a total of 43,855,652 single nucleotide polymorphisms (SNPs) after variant calling. We then filtered these SNPs using GATK VariantFiltration based on an examination of empirical distributions extracted using the GATK VariantsToTable function. We used the following filtering expression: ‘QUAL < 30 || MQ < 50.0 || SOR > 2.0 || QD < 10 || FS > 10.0 || MQRankSum < −1.0 || MQRankSum < 1.0 || ReadPosRankSum < −1.00 || ReadPosRankSum > 1.00,’ which applies stringent quality control to our variant calls. Specifically, this filtering removes variants with low Phred-scaled quality scores (QUAL < 30), low mapping quality (MQ < 50.0), high strand bias (SOR > 2.0), low confidence in the variant call normalized by read depth (QD < 10), significant strand bias detected by the Fisher test (FS > 10.0), and discrepancies in the mapping quality and read position between the reference and alternate alleles (MQRankSum and ReadPosRankSum). These criteria retain only high-confidence variants that are less likely to be affected by sequencing errors or biases for further analysis. After the GATK hard filtering, 26,892,855 biallelic SNPs remained. Further filtering with VCFtools v0.1.16 (Danecek et al., 2011. b) for a minimum quality of 30, a maximum of 5% missingness (retain only variants that have been successfully genotyped in at least 95% of individuals), and minor allele frequency (MAF > 0.05) resulted in a set of 1,577,126 SNPs. We further filtered this set of SNPs by linkage disequilibrium pruning (r2 > 0.1 within 10-kb windows) with bcftools v1.19 (Danecek et al., 2021.a) to result in a final dataset with 158,235 SNPs, which we used for population structure (DAPC, ADMIXTURE, pcadapt, sNMF, IBE/IBD), demographic inference (momi2), and all selection scans (pcadapt, pRDA).

### Genetic Diversity

*Genome-wide Heterozygosity—*To assess genome-wide (global) nuclear heterozygosity across both native and invasive populations, we employed ANGSD v0.933 (Korneliussen et al., 2014) as described in de Jager, D. (2021), which calculates heterozygosity from genotype likelihoods rather than called genotypes—an approach particularly advantageous for lower-coverage samples and genome wide heterozygosity patterns. For each deduplicated BAM file, we generated per-sample Site Frequency Spectra (SFS) using the ANGSD –dosaf command with the folded spectrum option (–fold 1). We filtered reads by removing those with mapping quality below 30, base quality below 20, improper pairing, non-unique mapping positions, and those flagged as low quality. We applied the Samtools genotype likelihood model (–GL 1), which accounts for sequencing errors and improper base calls. Individual heterozygosity was then calculated by using the realSFS program to extract the proportion of heterozygous sites from each sample’s SFS, dividing the number of heterozygous positions (second entry in the SFS) by the total number of sites (sum of first and second entries). We evaluated the robustness of heterozygosity comparisons between USA and China given uneven sample sizes (China: n=20, USA: n=98) using two complementary approaches. First, non-parametric bootstrap resampling (1,000 iterations) generated confidence intervals for mean differences while preserving original data structure. Second, random subsampling analysis repeatedly selected 20 USA individuals (matching China sample size) and performed balanced Welch’s t-tests across 1,000 iterations, recording p-values, effect sizes, and subsampled means to establish a null expectation for USA heterozygosity. We calculated the percentage of significant results (p < 0.05) to determine whether findings exceeded chance expectations (∼5%). This dual framework verifies that observed heterozygosity patterns reflect genuine invasion-associated genetic diversity loss rather than statistical bias from unbalanced sampling design.

*Runs of homozygosity and inbreeding coefficient—*Runs of homozygosity (ROH) were identified using PLINK v1.9 (Purcell et al., 2007) following recommendations by Meyermans et al. (2020) for non-model species. Only autosomal chromosomes were included in the analysis (excluding the sex chromosome: chr_4). Following the recommended approach for ROH detection in non-model organisms, we did not perform minor allele frequency pruning (––maf), or linkage disequilibrium (LD) pruning, as these filtering steps can lead to underestimation of true ROH segments in populations with complex demographic histories (Meyermans et al., 2020). Our input dataset consisted of 5,243,494 autosomal SNP variants across 118 individuals that passed our GATK hard filtering criteria and had a maximum missing rate of 5%, resulting in a high overall genotyping rate of 96.9%. We implemented a sliding window approach with a window size of 50 SNPs (––homozyg-window-snp 50) to detect homozygous segments. Within each window, we permitted at most 1 heterozygous SNP (––homozyg-window-het 1) and 5 missing genotypes (––homozyg-window-missing 5) to account for potential sequencing errors while preserving the integrity of homozygous segment detection. An SNP was classified as part of an ROH if it appeared in at least 5% of overlapping homozygous windows (––homozyg-window-threshold 0.05). To ensure robust ROH identification, we defined minimum criteria for ROH segments: each ROH required at least 50 consecutive SNPs (––homozyg-snp 50) and a minimum physical length of 500 kb (––homozyg-kb 500) to exclude short segments potentially resulting from linkage disequilibrium rather than identity by descent. We specified a maximum gap of 1000 kb between consecutive SNPs (––homozyg-gap 1000) to prevent fragmentation of ROH segments due to low marker density in certain genomic regions, while also setting a minimum density threshold of 1 SNP per 50 kb (––homozyg-density 50) to ensure sufficient marker coverage within detected ROH segments.

Following established protocols (Xu et al., 2019; Liu et al., 2022), ROHs were categorized into three size classes: small (0.5–1 Mb), medium (1–5 Mb), and large (>5 Mb). Summary statistics, including the number and total length of ROHs in each size class, were calculated for each individual and aggregated by geographic area to assess regional variations. The genomic inbreeding coefficient (FROH) was calculated by normalizing the total length of ROHs by the total length of the autosomal genome covered by SNPs (McQuillan et al., 2008), representing the proportion of the autosomal genome contained within homozygous segments above the specified threshold.

*Calculation of Coalescence Times—*The inverse relationship between ROH segment length and their approximate coalescence times is well established, with longer homozygous tracts typically representing more recent inbreeding events (Kardos et al., 2017). Following the mathematical framework described by Thompson (2013), we estimate the time to the most recent common ancestor (TMRCA) using the equation relating genetic length (L in cM) to generations (t) as L = 100/(2t). For each ROH size class in each population, we calculated the median physical length (Mb) and converted it to genetic distance (cM) using the empirically estimated recombination rate of 1 cM/Mb derived from our LD decay analysis to calibrate our runs of homozygosity (ROH) analysis and calculate time to TMRCA for different ROH length classes. For each ROH size class in each population (Shanghai and USA), we calculated the median physical length (in Mb) and converted it to genetic distance using the 1 cM/Mb recombination rate. The TMRCA in generations was then calculated as t = 100/(2L), and converted to years with one generation per year for the Spotted Lanternfly.

### Population Structure

*DAPC—*We analyzed population structure using Discriminant Analysis of Principal Components (DAPC) with adegenet (Jombart, 2008; Jombart & Ahmed, 2011) in R version 4.3.1 (R Core Team, 2023; RStudio Team, 2023), which identifies clusters of genetically related individuals while maximizing group separation. Cross-validation was used to select the optimal number of principal components, minimizing overfitting. The number of principal components with the lowest Mean Squared Error (MSE) was retained (Fig. S4).

*Ancestry Proportions—*To estimate individual ancestry proportions, we implemented the sparse non-negative matrix factorization (sNMF) approach using the LEA package in R (Frichot et al., 2014). We converted our filtered VCF file to the appropriate format for LEA using the vcftools2lfmm function. To determine the optimal number of ancestral populations (K), we tested values ranging from K=1 to K=10, with 10 independent runs for each value. For each run, the snmf function computes least-squares estimates of ancestry proportions and ancestral allele frequencies under the specified K value (Frichot et al., 2014). Model selection was performed using the cross-entropy criterion, which evaluates the prediction error of the model based on masked genotypes. Lower cross-entropy values indicate better predictive capabilities and thus a more appropriate value of K. For each K value, we calculated the mean cross-entropy across the 10 replicate runs to identify the most robust estimate of population structure. The run with the lowest cross-entropy for the optimal K value was selected for the final ancestry coefficient estimates (Fig. S5).

We also implemented the ADMIXTURE algorithm (Alexander et al., 2009), a model-based maximum likelihood clustering approach, to estimate individual ancestry proportions given a predefined number of ancestral populations (K). Analyses were conducted separately for the Shanghai and USA populations using genotype data converted into PLINK .bed format. We tested values of K ranging from 1 to 10, with 10 independent replicate runs per K using different random seeds to ensure convergence and robustness. Each run included fivefold cross-validation (––cv) to evaluate model predictive performance. Cross-validation (CV) error was extracted from each run, and the mean CV error for each K value was calculated across replicates. The optimal number of clusters was identified as the K value with the lowest average CV error (Fig. S5).

*Isolation by distance and environment—*We conducted isolation by distance (IBD) and isolation by environment (IBE) analyses. Genetic distances were calculated as 1 minus the identity-by-state (IBS) values using PLINK v1.09 (Purcell et al., 2007), providing a measure of genetic divergence, with values closer to 1 indicating greater dissimilarity.

We calculated geographic distances between the precise latitude and longitude for every sampled individual in the USA and between sampled sites in China. We quantified environmental variables at each location using remote-sensed data processed in Geographic Information Systems (GIS). We included bioclimatic variables from WorldClim2 (Fick & Hijmans, 2017), which included 11 temperature-related metrics (bio1-bio11) and 8 precipitation-related metrics (bio12-bio19). We also incorporated four urbanization metrics: tree canopy cover (TCC) from the Global 30m Landsat Tree Canopy Version 4 (Sexton et al., 2013), artificial light at night (ALAN) from VIIRS Plus DMSP Change in Lights dataset at 500m resolution (Small et al., 2020), human population density from Gridded Population of the World Version 4 at 1KM resolution (CIESIN, 2017), and artificial impervious surface (AIS) from the global impervious surface area dataset at 10 m resolution (Huang et al., 2022). To ensure consistent spatial resolution across all environmental layers, we resampled all 1 km resolution rasters (bioclimatic variables and population density) to 100 m resolution using bilinear resampling in ArcGIS Pro (Esri Inc., 2016), which is appropriate for continuous environmental variables. For each sampling location, we extracted the mean of each variable within a 300-meter buffer using ArcGIS Pro (Esri Inc., 2016). To reduce dimensionality, we performed separate principal component analyses (PCAs) on three categories of environmental variables: temperature (bio1-bio11), precipitation (bio12-bio19), and urbanization (ALAN, AIS, TCC, and population density). For each PCA, we extracted the first principal component (PC1) for use in subsequent analyses, as these captured the majority of variation (70%, 55%, and 68.4% for temperature, precipitation, and urbanization variables, respectively; Fig. S6). For each pair of sampled individuals, we calculated environmental distance as the Euclidean distance between their positions in this three-dimensional environmental PC space.

Both simple and partial Mantel tests were performed using the ecodist package (Goslee & Urban, 2007) with 1,000 permutations to test for correlations between genetic distances and geographic distances (IBD), as well as between genetic distances and environmental distances (IBE). Additionally, we performed a partial Mantel test to assess the correlation between genetic and environmental distances while controlling for geographic distance.

### Demographic History

*LD Decay—*Linkage disequilibrium (LD) decay was assessed by PopLDdecay (v3.43) to calculate LD decay patterns across different populations (Zhang et al. 2018). The analysis was performed with a maximum distance of 300 kb between SNP pairs. We used the dataset that has 1,577,126 SNPs without LD pruning, as described above. LD decay curves were generated by plotting the squared correlation coefficient (r²) between SNP pairs against their physical distance in kilobases using *Plot MultiPop.pl* script provided by PopLDdecay (Zhang et al. 2018).

To estimate population-specific recombination rates from LD decay patterns, we implemented a nonlinear regression approach following established empirical methods (Remington et al., 2001; Weiss & Clark, 2002). While Sved (1971) provides the theoretical expectation E[r²] = 1/(1 + 4Nₑcρ), empirical studies commonly use exponential decay models for robust parameter estimation across diverse genomic contexts (Hill & Weir, 1988; Marroni et al., 2011).

We modeled LD decay using: r² = b + (d₀ – b) × exp(–c × d), where r² represents the squared correlation coefficient between alleles at two loci; d is the physical distance between loci in base pairs; d₀ is the theoretical initial r² at zero distance (representing maximum possible LD); b is the asymptotic r² at infinite distance (background LD); and c is the decay rate parameter that determines how rapidly LD decreases with physical distance. The parameter c was scaled by 100,000 to provide units comparable to centimorgans per megabase for comparative purposes between populations. To ensure robust parameter estimation, we implemented a grid search approach using the nls2 package in R (Grothendieck G, Team RC 2024). We explored a wide range of biologically plausible parameter values: initial r² (d₀) from 0.8 to 1.0, asymptotic r² (b) from 0.1 to 0.3, and rate parameters (c) from 0.000001 to 0.0003, corresponding to recombination rates of approximately 0.1 to 30 cM/Mb. This comprehensive parameter space covered the full range of recombination rates reported in insects, from species with low rates (e.g., some *Lepidoptera* at ∼0.5 cM/Mb) to those with high rates (e.g., *Hymenoptera* at >10 cM/Mb). Population-specific recombination rates were estimated separately for Shanghai and USA populations to detect any potential differences in recombination landscapes between native and invasive populations. Model fitting was performed using the brute-force algorithm in nls2 to thoroughly explore the parameter space and avoid local optima. The best-fitting model for each population was selected based on minimizing the residual sum of squares.

*Nuclear Phylogenetic inference—*We inferred a maximum likelihood tree of genetic relationships among individuals. We prepared the filtered VCF file containing all 118 samples using VCFtools to retain only genotype information, then converted it to PHYLIP format using the vcf2phylip.py script (Ortiz, 2019; Danecek et al., 2021.a). We performed phylogenetic inference using IQ-TREE v1.6.12 (Nguyen et al., 2015) with ModelFinder (Kalyaanamoorthy et al., 2017) to determine the optimal evolutionary model. For the complete dataset of all samples, ModelFinder tested 242 DNA models with a sample size of 158,123 SNPs. Based on the Bayesian Information Criterion (BIC), the GTR+F+ASC+G4 model was identified as the best-fit model. We used 1000 bootstrap replicates implemented with the SH-aLRT test (–bb 1000) to assess branch support and specified DNA sequence data (–st DNA) with the selected model (–m GTR+F+ASC+G4). The resulting maximum-likelihood tree was midpoint-rooted and visualized using the phytools package in R (Revell, 2012).

*Mitochondrial haplotype analysis—*Illumina paired-end reads were trimmed and quality-filtered prior to alignment. Reads were mapped to the *Lycorma delicatula* mitochondrial reference genome (GenBank accession EU909203.1) using BWA-MEM v0.7.17 (Li, 2013). Alignments were sorted and converted using SAMtools v1.14 (Danecek et al., 2021). Read groups were added with Picard Tools v2.17.11 (Broad Institute, 2019), and duplicates were marked using GATK MarkDuplicates from GATK v4.3.0.0 (Van der Auwera & O’Connor, 2020). Variant calling was performed using GATK HaplotypeCaller in GVCF mode with –-sample-ploidy 1 to reflect mitochondrial haploidy. Joint genotyping across all samples was conducted with GATK’s GenotypeGVCFs. SNPs were extracted using GATK SelectVariants and filtered with VariantFiltration using the following thresholds: QUAL < 30.0, QD < 2.0, MQ < 40.0, FS > 60.0, SOR > 3.0, MQRankSum < –12.5 or > 12.5, ReadPosRankSum < –8.0 or > 8.0. PASS-filtered variants were used to reconstruct full mitochondrial consensus sequences for each individual using bcftools consensus (Li et al., 2009). To generate high-confidence individual mitochondrial consensus sequences, we applied a site-level allelic support threshold of 75%— consistent with the default consensus calling criteria in Geneious and following the approach of Du et al. (2021). We then integrated our 118 consensus mitogenomes with 392 previously published Spotted Lanternflies mitochondrial genomes from Du et al. (2021), resulting in a combined dataset of 510 sequences. These were aligned using MAFFT v7.475 with the high-accuracy G-INS-i strategy (––globalpair –-maxiterate 1000) to perform global alignment optimized for full-length sequences (Katoh & Standley, 2013). The alignment was manually inspected for sequence orientation and length uniformity. We identified 446 unique haplotypes from the 510 complete mitochondrial genomes using the pegas R package (Paradis, 2010). We inferred a haplotype phylogeny using IQ-TREE v1.6.0 (Nguyen et al., 2015), with the mitogenome of *Lycorma meliae* (MT079725.1) included as an outgroup to root the tree. The best-fit substitution model was selected using ModelFinder Plus, which identified TIM+F+R3 as the optimal model based on the Bayesian Information Criterion (Kalyaanamoorthy et al., 2017). Node support was assessed with 1,000 ultrafast bootstrapreplicates and SH-aLRT tests. We then used ggtree (Yu, 2020) to plot the unrooted tree of the .contree from the IQ-TREE result and assigned haplotype groups based on monophyletic clades aligning with the haplotypes assigned by Du et al. (2021).

*Momi2 demographic modelling—*We inferred demographic history using momi2 (Moran Models for Inference), a Python package designed for constructing and comparing empirical and simulated demographic models (Kamm et al., 2022). Momi2 is particularly effective in evaluating recent demographic changes using whole genomes from contemporary populations, providing robust inferences by accurately reconstructing population size dynamics (Reid & Pinsky, 2022). Given the recent invasion of Spotted Lanternflies in the United States, this approach is appropriate to model more recent demographic changes, such as within 100 years. Other methods, such as PSMC, focus on much older events, modeling ancient demographic history from hundreds of thousands to millions of years ago, offering limited insights into the recent demographic history of invasions. To prepare the dataset for momi2 analysis, we split the main VCF file into population-specific VCFs using bcftools v1.14 (Danecek et al., 2021) and generated corresponding BED files to represent genomic regions. Allele count files for each population were created using the momi.read_vcf function in momi2. The site-frequency spectrum (SFS) was extracted from these allele counts using the momi.extract_sfs function.

We built eight models representing different historical scenarios, each designed based on plausible historical events from literature (Du et al., 2021; Kim et al., 2021). These models allowed us to test hypotheses about recent demographic changes but not make definitive claims about a specific invasion route. The models constructed are as follows:

*Model 1 Urban Bridgehead*: Shanghai rural → Shanghai urban → South Korea → USA.

*Model 2 Urban Direct*: Shanghai rural → Shanghai urban → USA.

*Model 3 Rural Bridgehead*: Shanghai rural → South Korea → USA (with separate Shanghai rural → Shanghai urban).

*Model 4 Rural Direct*: Shanghai rural → USA (with separate Shanghai rural → Shanghai urban).

*Model 5 Independent Urban Bridgehead*: Common Ancestral → Shanghai rural/urban (parallel), Shanghai urban → South Korea → USA.

*Model 6 Independent Urban Direct*: Common Ancestral → Shanghai rural/urban (parallel), Shanghai urban → USA.

*Model 7 Independent Rural Bridgehead*: Common Ancestral → Shanghai rural/urban (parallel), Shanghai rural → South Korea → USA.

*Model 8 Independent Rural Direct*: Common Ancestral → Shanghai rural/urban (parallel), Shanghai rural → USA.

Topologies for the demographic models described above are illustrated in Fig. S7. Details of all parameters used in each of the eight demographic models, including population sizes and the timing of each demographic event, can be found in the supplementary materials (Table S1). All models were optimized using the L-BFGS-B algorithm with 200 iterations and convergence tolerances (ftol and gtol) of 1 × 10^-6. For each model, we performed 785 bootstrap replicates by resampling the SFS to estimate confidence intervals for all parameters. Model selection was based on the Akaike Information Criterion (AIC), delta AIC, and AIC weight. The Akaike Information Criterion (AIC) is a statistical measure used for model selection that balances model fit against complexity. It is calculated as AIC = 2k – 2ln(L), where k is the number of estimated parameters and L is the maximum likelihood of the model. This formula penalizes models with more parameters to prevent overfitting, as each additional parameter must provide sufficient improvement in likelihood to justify its inclusion. Delta AIC (ΔAIC) represents the difference between each model’s AIC value and the lowest AIC value among all candidate models: ΔAIC = AICi – AICmin. AIC weight (wi) quantifies the relative probability that a given model is the best among the candidate set, calculated as wi = exp(–0.5 × ΔAICi) / Σ exp(–0.5 × ΔAICj). The optimal model was identified as having the lowest AIC value and highest AIC weight for each run. For the best-fitting model, we calculated 95% confidence intervals for all parameters based on the bootstrap distribution. In all models, we do not specify whether the demographic change happens gradually or abruptly; rather, the model estimates population size changes and split events.

### Signature of Selection

*SNP annotation—*We annotated the VCF files using SnpEff v4.3t (Cingolani et al., 2012) to predict and classify the functional effects of identified variants. First, we constructed a custom SnpEff database for the Spotted Lanternfly reference genome (Snead et al. 2025) using the genomic sequence (SLF_Hap1.fasta), gene feature annotations (SLF_Hap1.gff3), and protein sequence data (Hap1.faa). The database was configured to recognize the Spotted Lanternfly genome architecture and gene structure definitions. Each variant was classified according to its genomic location (e.g., intergenic, intro_variant, up_stream or down_stream of a gene coding region, etc.). This annotation allowed us to identify variants with potential functional significance. We only focus on downstream functional interpretation of genes with established functional annotations, excluding interpretation of SNPs in intergenic regions.

*PCAdapt*—We performed PCA-based genome scans using pcadapt v4.3.3 (Privé, F et al., 2020). We conducted separate analyses for Shanghai and USA populations: (1) to detect selection occurring in each geographic region and (2) to minimize the confounding effects of strong population structure between Shanghai and USA populations, which could otherwise mask within-region selection. For Shanghai populations, we selected K = 2 for subsequent analyses based on visual inspection of the scree plot and proportion of genetic variance captured in each PC (Fig. S8). In contrast, the USA populations exhibited weak structure and gradual decay in eigenvalues, consistent with a recent invasion and high gene flow. Due to the absence of a clear “elbow” in the scree plot (Fig. S8), we adopted a conservative multi-K strategy, running *pcadapt* with K = 2 through 6. At each K, candidate SNPs were identified using a q-value-based false discovery rate (FDR) threshold of < 0.01. FDR correction was applied using the qvalue function to control for multiple testing, with candidate loci defined as those with false discovery rate (FDR) adjusted p-values below α = 0.01, using Mahalanobis distance computed from the top K PCs. For the USA analysis, we then focused on conserved outliers—SNPs consistently identified as candidates across all five K values—which minimizes the risk of false positives due to noise or overfitting at any single K.

We performed separate Gene Ontology (GO) enrichment analyses for the candidate loci identified in the Shanghai and USA populations. For each population, genes containing candidate SNPs were tested against the complete set of annotated genes in the Spotted Lanternfly genome (Snead et al., 2025) using the goseq R package (Young et al., 2010). Enrichment was assessed independently for each region, with Benjamini–Hochberg correction applied to control for multiple testing. Significantly enriched GO terms were identified at a false discovery rate (FDR) threshold of 0.05.

*Population Branch Excess (PBE) Analysis—*To investigate population-specific signatures of selection in the native range of the Spotted Lanternfly, we conducted a Population Branch Excess (PBE) Analysis using a subset of 760 SNPs previously identified as outliers under selection (FDR < 0.01) in Shanghai populations by pcadapt. We first used VCFtools (Danecek et al., 2011) to calculate pairwise Weir and Cockerham FST values between three population pairs (Shanghai rural vs. urban, Shanghai rural vs. USA, and Shanghai urban vs. USA) using VCFtools. These FST values were transformed into divergence measures (T = –log(1 – FST)) to estimate branch lengths and compute standard PBS values following Yi et al. (2010). We performed all subsequent analyses in R. SNPs with missing values or FST values equal to or greater than 1 were removed, yielding a high-confidence set of 628 loci for downstream analyses. We transformed the pairwise FST values into divergence measures (T) using the equation T = –log(1 – FST). Standard PBS values for the Shanghai urban and rural populations were calculated following Yi et al. (2010) using the equation PBSA=(TAB+TAC−TBC)/2, where A is the focal population and B and C represent the other two populations. To account for demographic biases inherent in standard PBS calculations, we additionally computed Population Branch Excess (PBE) as described by Yassin et al. (2016).

The PBE for each population was calculated using the formula:

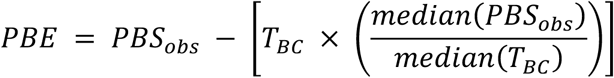

PBE specifically corrects for demographic biases by incorporating a normalization factor based on median values across loci (Yassin et al., 2016). We calculated PBE separately for each focal population to directly test for environment-specific selection. For the urban population, we computed PBE_Urban = PBS_Urban – (T_Rural_US × med_PBS_Urban/med_T_Rural_US), using the rural and US populations as references. Similarly, for the rural population, we calculated PBE_Rural = PBS_Rural – (T_Urban_US × med_PBS_Rural/med_T_Urban_US), using the urban and US populations as references. In each case, positive PBE values indicate stronger-than-expected selection in that specific environment after accounting for the shared demographic history between populations. This dual calculation allows for the independent identification of selection signals in each environment without assuming that selection in one environment necessarily corresponds to the absence of selection in the other. We visualized these environment-specific selection signals using mirrored Manhattan plots, emphasizing the most significant SNPs falling above PBE=0.1.

*Allele frequencies—*Allele frequencies for the 760 candidate SNPs identified by genome-wide selection scans were calculated for each population (Shanghai urban, Shanghai rural, and USA invasive populations) using VCFtools v0.1.16 (––freq option). Then we extracted alternative (ALT) allele frequencies, merged the datasets by SNP identifiers (CHROM and POS), and annotated them using gene annotation data. We calculated mean allele frequencies and standard errors for Shanghai urban, Shanghai rural, and USA populations, visualizing these differences using pheatmap in R (Kolde R. 2025).

*Partial RDA—*To identify genomic variants associated with environmental variation in USA Spotted Lanternfly populations, we performed redundancy analysis (RDA) using the R package vegan (Oksanen et al., 2022). We used the same environmental principal components described in section 2.4.2, incorporating the first principal component from each category (temperature, precipitation, and urbanization), confirming these predictors had variance inflation factors below 2. We performed partial RDA that accounted for spatial autocorrelation, allowing us to identify genetic variations associated with environmental adaptation while accounting for geographic proximity. We used distance-based Moran’s eigenvector maps (dbMEMs) constructed from a Gabriel graph neighborhood network with inverse distance weighting (km⁻¹) following Forester et al. (2018). Significant spatial components (MEM43, MEM53, MEM59, MEM64) were identified through forward selection (α = 0.05) using the forward.sel function (Dray et al., 2022) and incorporated into the partial RDA model. The statistical significance of the whole model and each RDA axis and predictor was assessed using 1,000 permutations. Candidate SNPs potentially under selection were identified as those with loadings exceeding out from the mean on significant RDA axes, following the approach established by Capblancq et al. (2018).

*Candidate SNPs—*To identify genomic regions under selection, we compared candidate SNPs identified by two approaches: partial redundancy analysis (pRDA) and PCAdapt. The overlapping SNPs were identified using the intersect function in R. We performed Gene Ontology (GO) enrichment analysis on the 141 SNPs identified by both selection methods using the goseq v1.50.0 (Young et al., 2010). Out of 184 SNPs, 158 were successfully mapped to annotated genes. Significantly enriched GO terms were identified using a threshold of FDR < 0.05 to account for the relatively small gene set. We calculated enrichment ratios (observed genes in category / expected genes in category) and examined the primary environmental predictor (temperature, precipitation, or urbanization) associated with each shared SNP based on the partial RDA results.

We analyzed heterozygosity patterns at these 184 candidate SNPs. Individual heterozygosity of these 184 SNPs was calculated using the –-het option in VCFtools, which measures observed and expected autosomal homozygous genotype counts for each sample. We then compared heterozygosity of these sites and allele frequencies across three populations: Shanghai rural (SH rural), Shanghai urban (SH urban), and USA populations.

### ANGSD/ngstools Comparison

To account for the moderate sequencing depth (3-6x coverage) of our dataset, we repeated our population genetics analysis using genotype likelihoods with ANGSD v0.933 (Korneliussen et al., 2014). This approach is recommended for low-to-medium coverage data as it incorporates uncertainty in genotype calls. We generated genotype likelihoods from deduplicated BAM files generated above using stringent filters (minimum mapping quality 30, base quality 20, minimum minor allele frequency 0.05, max missing 0.05, and SNP p-value cutoff 1×10⁻⁶), resulting in 6,183,248 variant sites across 118 individuals. To minimize the effects of linkage disequilibrium, we ran ngsLD within 10-kb windows and pruned sites with Pearson correlation coefficients (r²) ≥ 0.1 using the prune_graph tool, yielding 615,307 unlinked SNPs (Fox et al., 2019). Basically, we kept all filtering criteria the same as we did above for called genotypes.

Population structure was evaluated with PCAngsd v0.99 (Meisner & Albrechtsen, 2018), which computes principal components directly from genotype likelihoods, as well as via a phylogenetic analysis. Notably, the ANGSD-based results revealed population structure patterns highly consistent with those obtained from the GATK-called genotypes, indicating that our stringent filtering pipeline yielded robust and comparable SNP datasets across both approaches. This methodological cross-validation reinforces the reliability of our findings (Fig. S11).

We also computed pairwise identity-by-state (IBS) genetic distances using ANGSD v0.933 (–doIBS 1) using the above unlinked SNPs and repeated the isolation-by-distance (IBD) and isolation-by-environment (IBE) analyses using these genotype likelihood-derived distances. The IBD and IBE results based on ANGSD distances closely mirrored those obtained from hard genotype calls, further supporting the consistency and robustness of our analysis (Fig. S12).

## Acknowledgments

We acknowledge Tuc H.M. Nguyen for his invaluable assistance in setting up and sharing essential scripts for the GATK pipeline. His advice and support on mitochondrial genome assembly and haplotype analysis. We also extend our gratitude to the individuals who contributed to the sampling efforts for this project: Gaia Rueda Moreno, Hannah Owen, Karin Kiontke, Rafael Baez, Emerald Lin, Katie Schneider Paolantonio, Jake Lively, Ishani Sinha, Andrea Rummel, Tim Martin, Zulekha Munshi-South, Jonathan Zschau, Lee vonKraus, and Cleo Falvey. This work was supported in part through the NYU IT High-Performance Computing resources, services, and staff expertise. We acknowledge the Zegar Family Foundation for their generous support. We thank the NYU Center for Genomics and System Biology Genomics Core for their assistance and resources. A.A.S. was supported by NSF-PRFB (NSF-PRFB #2305939).

## Author Contributions

All authors conceptualized and designed the research. Y.Z. sampled the Spotted Lanternflies in the native range and performed DNA extraction in China. F.M. conducted the sequencing related benwork in the United States and performed all the downstream analyses. K.M.W., A.A.S., and J.M.S. provided assistance and troubleshooting with analyses. F.M. and K.M.W. drafted the manuscript. F.M., K.M.W., and A.A.S. arranged the figures. All authors (F.M., K.M.W., A.A.S., Y.Z., and J.M.S.) contributed to the writing of the manuscript and have approved the final version.

