## Supplementary Data for "Cities as evolutionary incubators for the global spread of the Spotted Lanternfly"

### Table of Contents:

Fig. S1: Sampling map
Fig. S2: Distribution of long ROHs across the genome in the USA invasive population
Fig. S3: Mitochondrial haplotype analysis
Fig. S4: DAPC: PCA and CV
Fig. S5: sNMF and ADMIXTURE analysis
Fig. S6: Environmental PCA analysis
Fig. S7: Topologies for the Momi2 demographic models tested
Fig. S8: PCAdapt scree plot for SH and USA
Fig. S9: LD patterns across chromosomes of native Shanghai population
Fig. S10: Observed heterozygosity (Ho) at 184 candidate SNPs across populations
Fig. S11: Population Structure using ANGSD/ngstools
Fig. S12: Isolation-by-distance (IBD) and isolation-by-environment (IBE) analyses based on pairwise genetic distances calculated using genotype likelihoods in ANGSD

Table S1: Estimated time to most recent common ancestor (TMRCA) from ROHs
Table S2: Momi2 parameters
Table S3: Momi2 AIC statistics for model selection
Table S4: Momi2 best model statistics
Table S5: SNP density across chromosomes in Spotted Lanternflies of the USA population

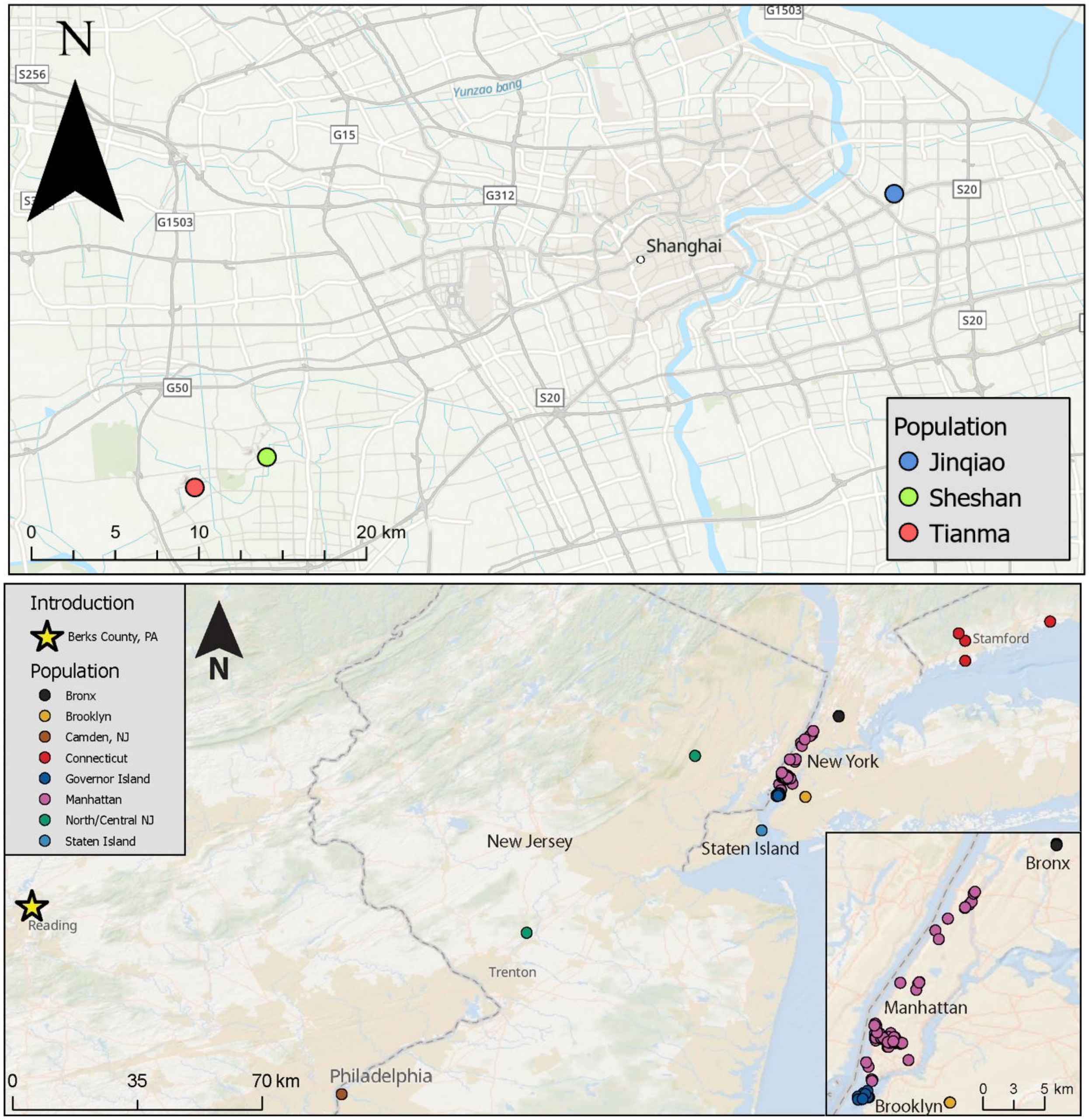


Fig. S1. Geographic distribution of Spotted Lanternfly sampling locations in the native and invasive ranges. Maps show the locations of spotted lanternfly collections from the native range (Shanghai, China; upper panel) and the invasive range in the northeastern United States (lower panel). Each point represents a unique sampling site, with colors corresponding to different locations.

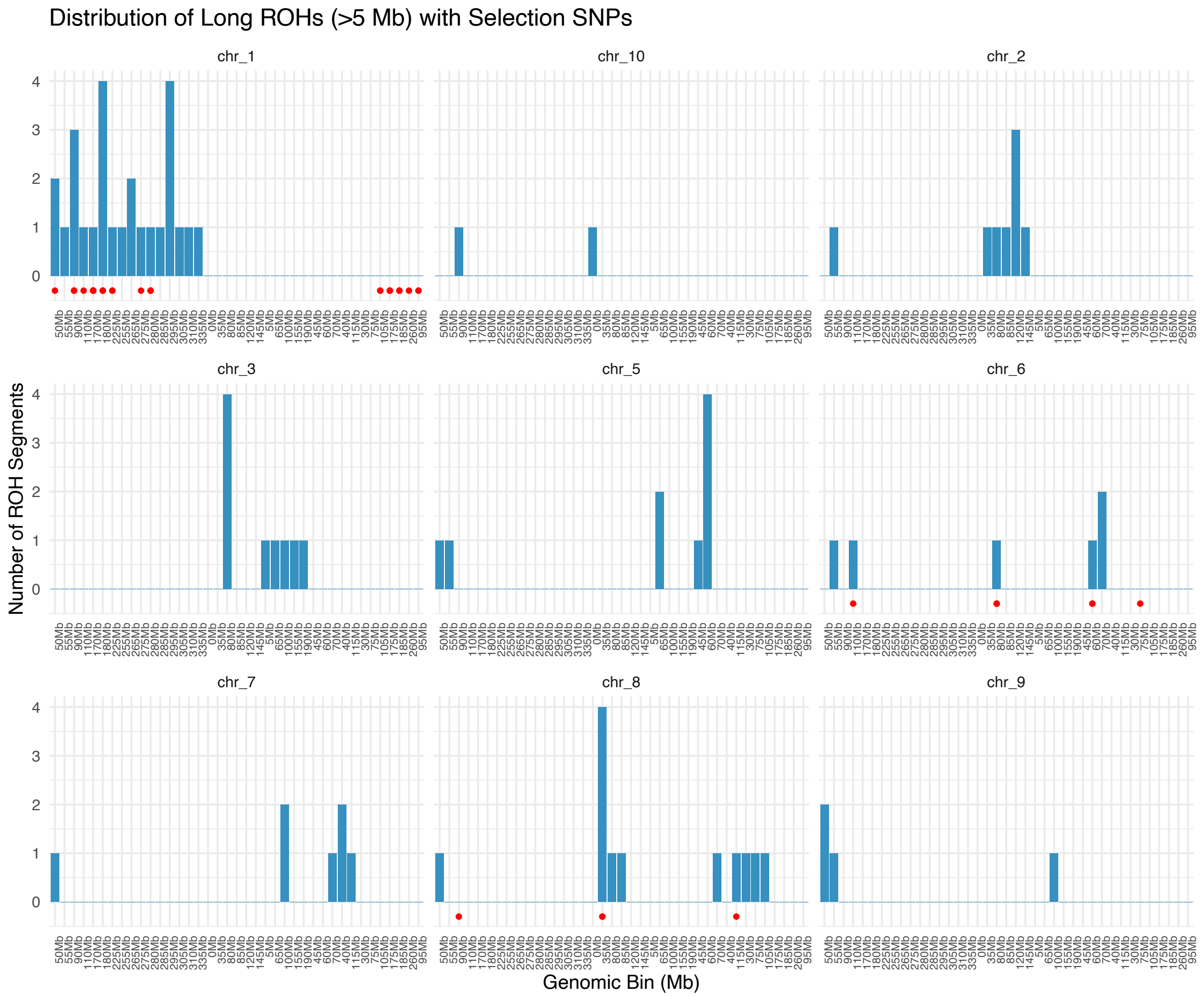


Fig. S2. Distribution of long runs of homozygosity (ROHs) across the genome in the USA invasive population. Each panel shows the number of long ROH segments (>5 Mb) per 5 Mb non-overlapping bin across chromosomes 1–13. Red dots indicate the genomic positions of SNPs identified as selection candidates in the USA population, which overlap with long ROH segments.

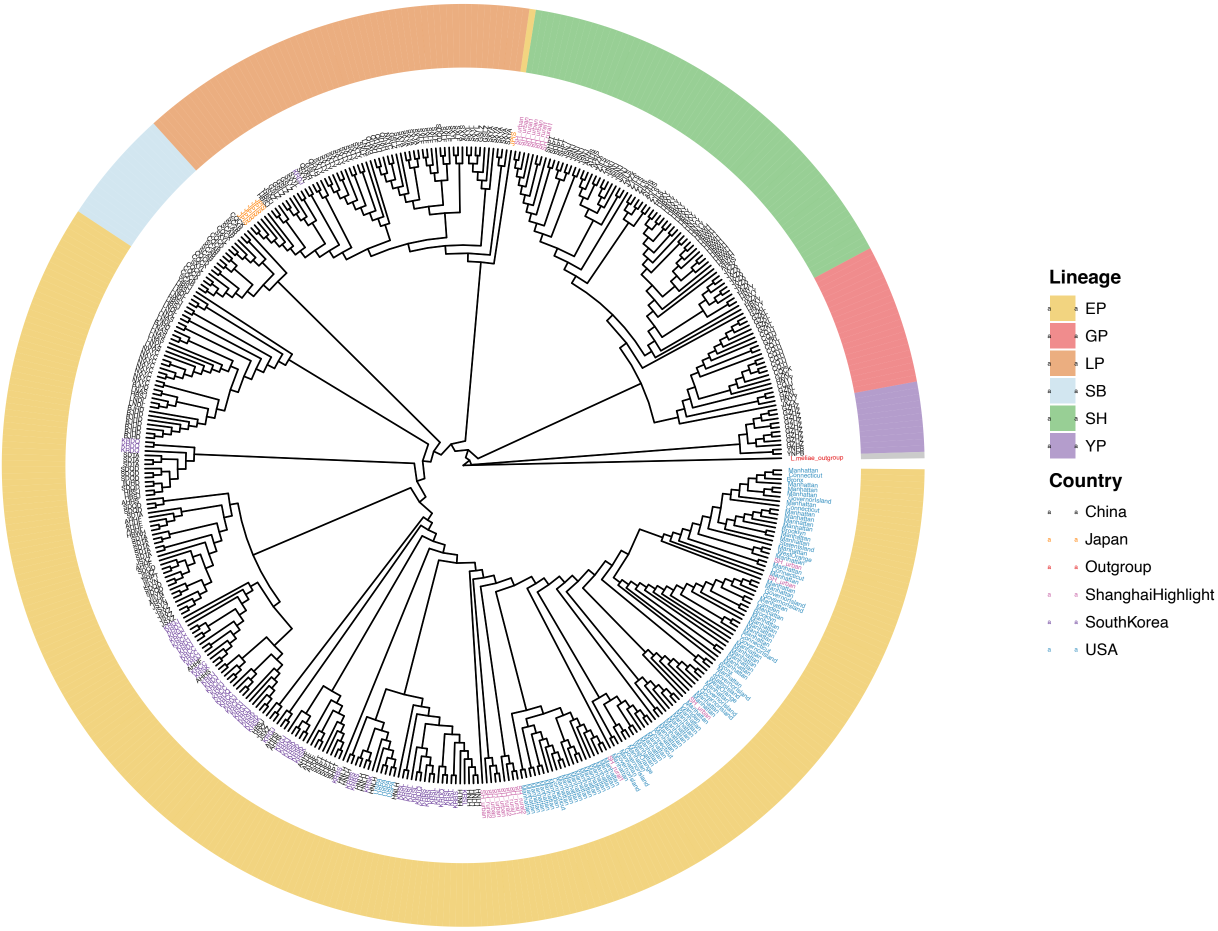


Figure S3. Maximum likelihood phylogeny of mitochondrial haplotypes from 511 individuals, including 118 newly sequenced in this study (shown in bold italic) and 392 previously published by Du et al. (2021), with *Lycorma meliae* as outgroup. Tip points are colored by country, and samples from this study collected in Shanghai are additionally highlighted in pink. The colored outer ring indicates the six major mitochondrial haplogroups defined by Du et al. (2021): East Plain (EP), Southeast Hills (SH), Sichuan Basin (SB), Yunnan Plateau (YP), Guizhou Plateau (GP), and Loess Plateau (LP). The invasive U.S. samples exclusively cluster within the EP lineage, reaffirming this region as the most likely source of the U.S. invasion. Both EP and SH haplotypes were found in urban and rural Shanghai locations sampled.

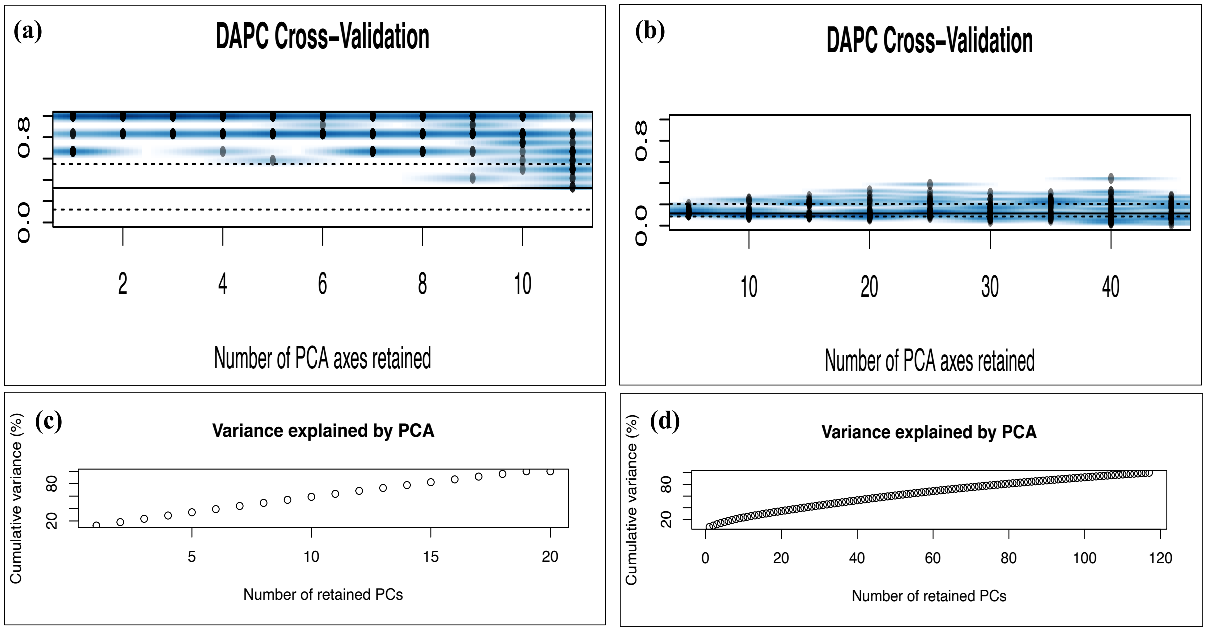


Fig. S4. Cross-validation analysis for determining optimal parameters in DAPC for spotted lanternfly populations. (a) For Shanghai populations (native range), cross-validation indicates optimal performance with just 2 principal components (PCs), achieving 97.2% successful assignment accuracy with lowest mean squared error (MSE). (b) For USA populations (invasive range), cross-validation supports retaining 35 PCs with substantially lower assignment accuracy (17.0%), indicating much weaker population structure. (c) Variance explained by retained PCs in the Shanghai dataset, where just 2 PCs capture sufficient genetic variation to achieve high discrimination between urban and rural populations. (d) Variance explained by retained PCs in the USA dataset, where 35 PCs were needed to capture significant genetic variation.

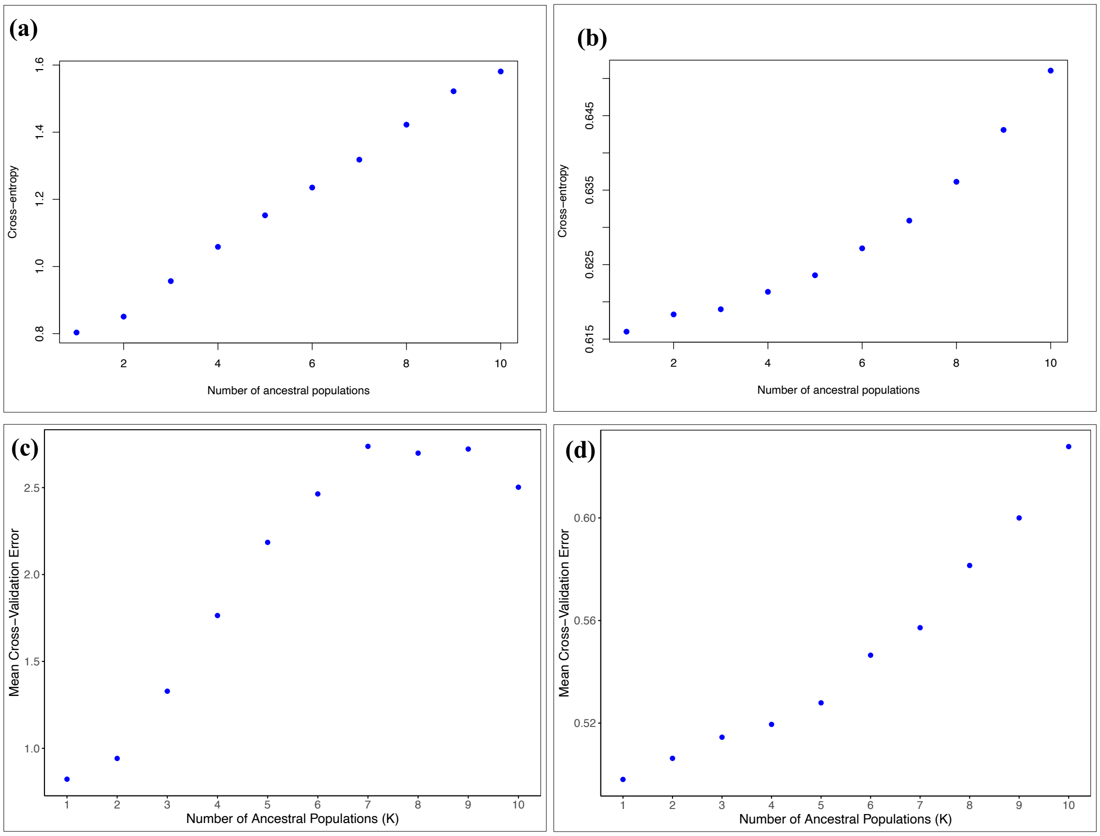


Fig. S5. Comparison of population structure inference using sNMF and ADMIXTURE. Both methods support minimal genetic structure (K = 1) for both Shanghai and USA populations. Panels show mean cross-entropy or CV error values across K = 1 to 10: **(a)** sNMF cross-entropy for Shanghai. **(b)** sNMF cross-entropy for USA. **(c)** ADMIXTURE cross-validation error for Shanghai. **(d)** ADMIXTURE cross-validation error for USA.

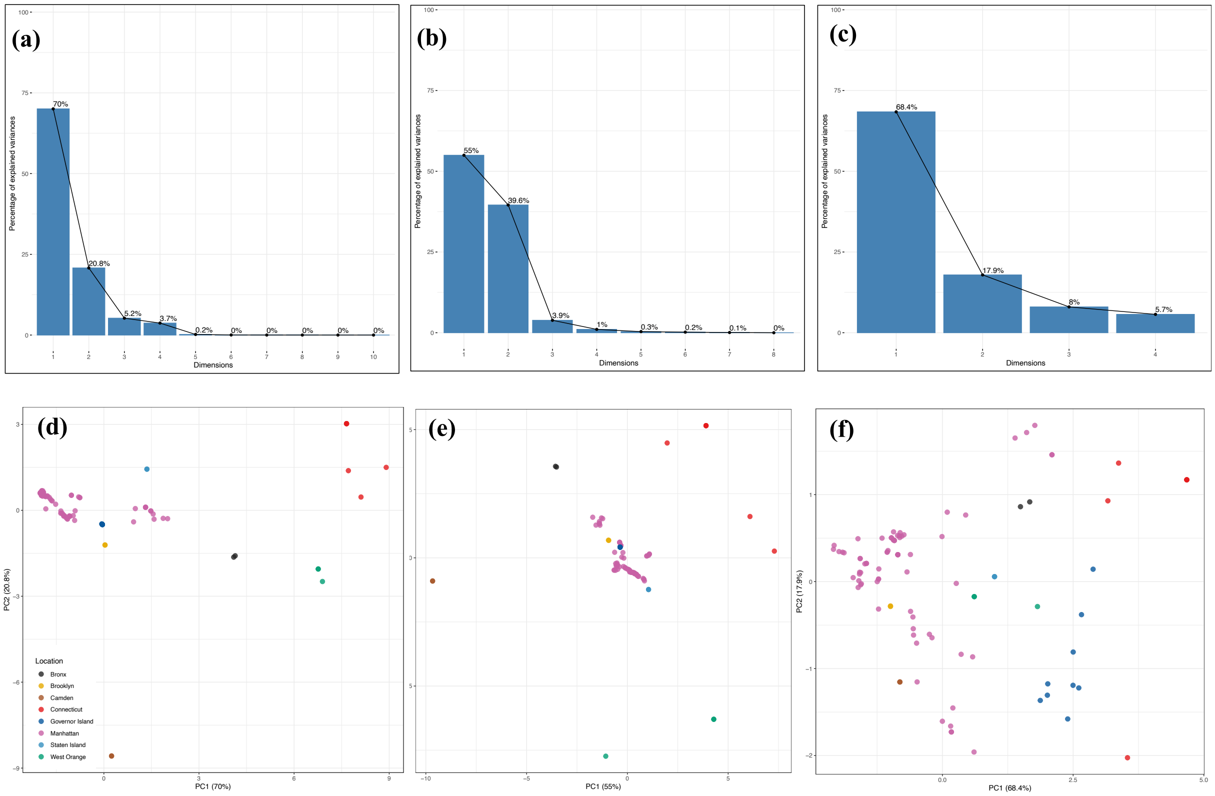


Fig. S6. Principal Component Analyses (PCAs) of environmental variables across sampling locations in the USA. Top row displays scree plots showing the percentage of variance explained by each principal component for (a) temperature variables (bioclimatic variables 1-11), (b) precipitation variables (bioclimatic variables 12-19), and (c) urbanization metrics (artificial light at night, impervious surface, tree canopy cover, and population density). Bottom row shows biplots of the first two principal components for (d) temperature, (e) precipitation, and (f) urbanization variables, with points colored by sampling location. The first two principal components capture 90.8%, 94.5%, and 86.3% of the total variation for temperature, precipitation, and urbanization, respectively. Temperature PC1 (70.1% variance) contrasts stable, warm environments (negative values) with environments exhibiting high temperature fluctuations (positive values). Precipitation PC1 (55.0% variance) distinguishes consistently wet environments (positive values) from those with stronger wet/dry seasonality (negative values). Urbanization PC1 (68.4% variance) primarily reflects a gradient from highly urbanized to more natural areas.

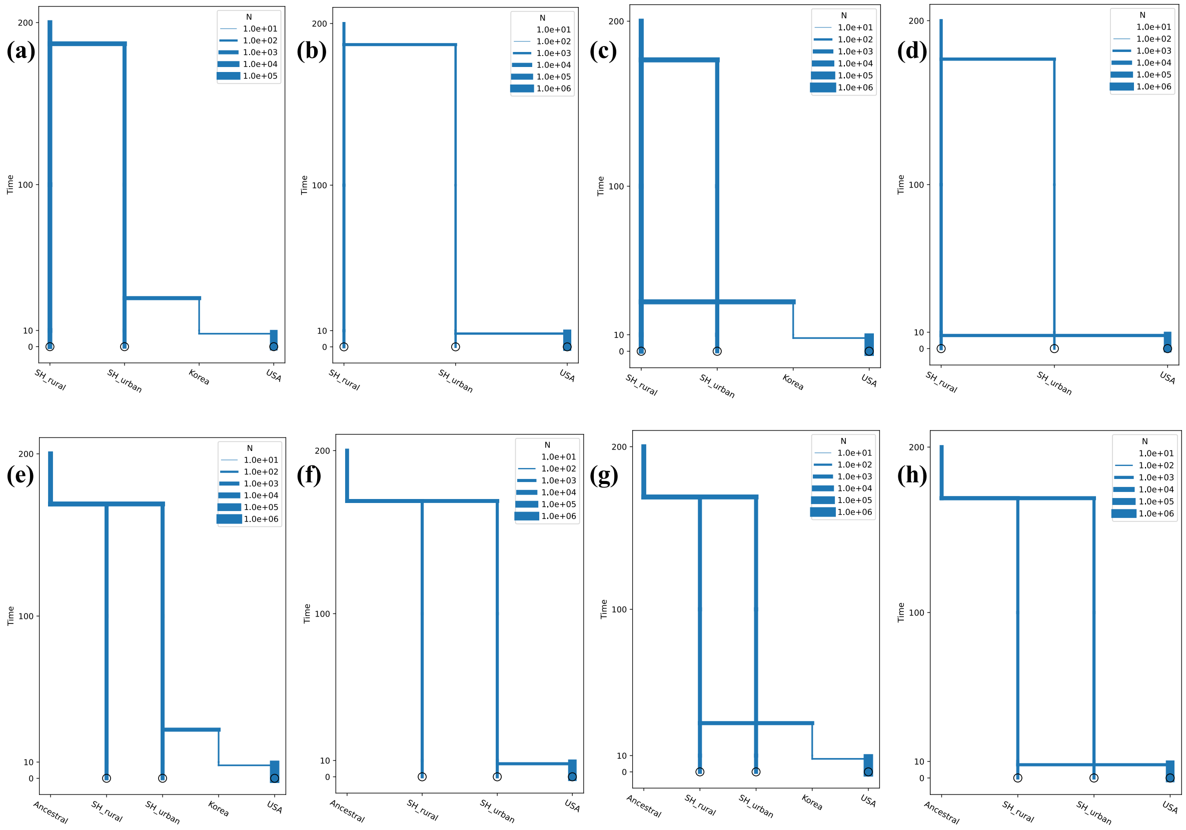


Fig. S7.Topologies for the demographic models tested. Models differ in their source location in China (rural vs. urban Shanghai), route of invasion (direct vs. bridgehead through South Korea), and evolutionary relationship between Shanghai populations (sequential vs. independent). (a) Model 1 (Bridgehead Model): Sequential urbanization in Shanghai followed by introduction to South Korea and subsequent colonization of the USA; (b) Model 2 (Direct Invasion Model): Sequential urbanization in Shanghai followed by direct introduction to the USA; (c) Model 3: Rural origin with Korea as bridgehead, where Shanghai rural populations are ancestral to both Korean and Shanghai urban populations; (d) Model 4: Direct rural origin with parallel urbanization in Shanghai; (e) Model 5: Independent evolution of Shanghai urban and rural populations with urban-sourced bridgehead through Korea; (f) Model 6: Independent evolution of Shanghai urban and rural populations with direct urban-sourced invasion; (g) Model 7: Independent evolution of Shanghai urban and rural populations with rural-sourced bridgehead through Korea; (h) Model 8: Independent evolution of Shanghai urban and rural populations with direct rural-sourced invasion. Time is shown on the y-axis (not to scale) and effective population size (N) is represented by varying branch widths.

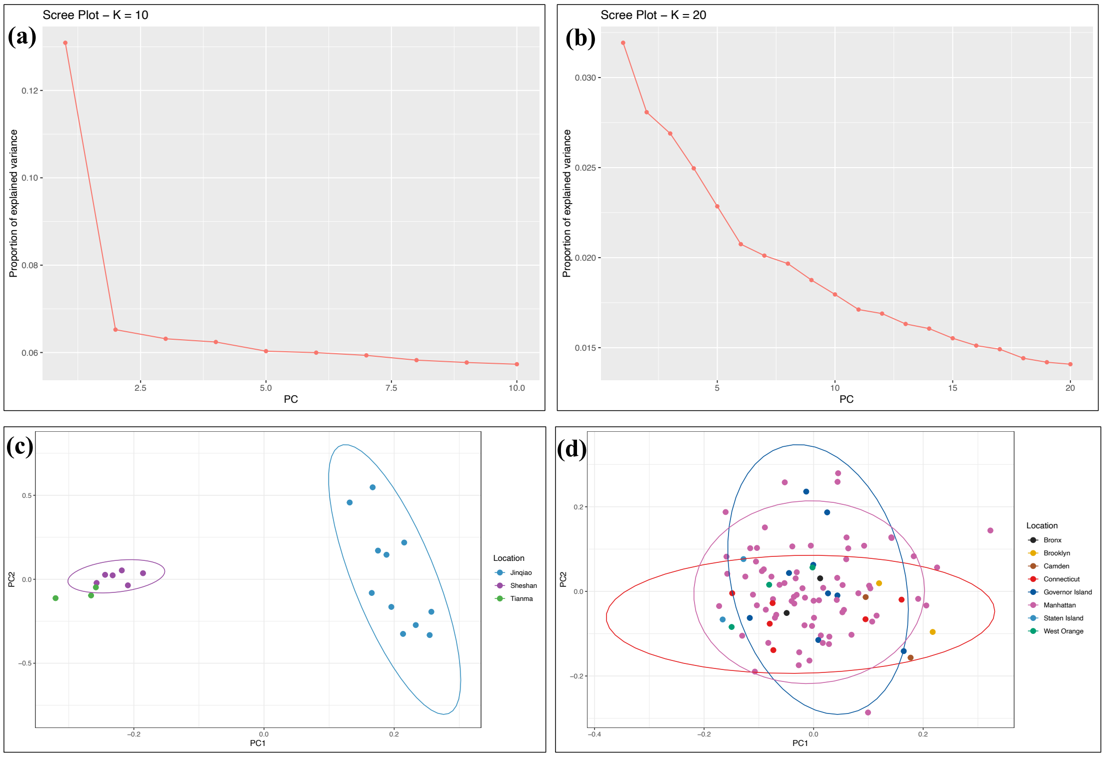


Fig. S8. PCAdapt for native and invasive spotted lanternfly populations. (a) Scree plot for Shanghai (native) locations with K=10 principal components, showing a sharp decline in explained variance after the first two components. (b) Scree plot for USA (invasive) locations with K=20, displaying a gradual decline in explained variance without a clear "elbow" point, typical of recently invaded populations with limited genetic differentiation. (c) Population structure visualization of Shanghai samples along PC1 and PC2, revealing clear separation between urban (blue) and rural (green and purple) populations. (d) Population structure visualization of USA samples, showing extensive overlap among sampling locations spanning the northeastern United States.

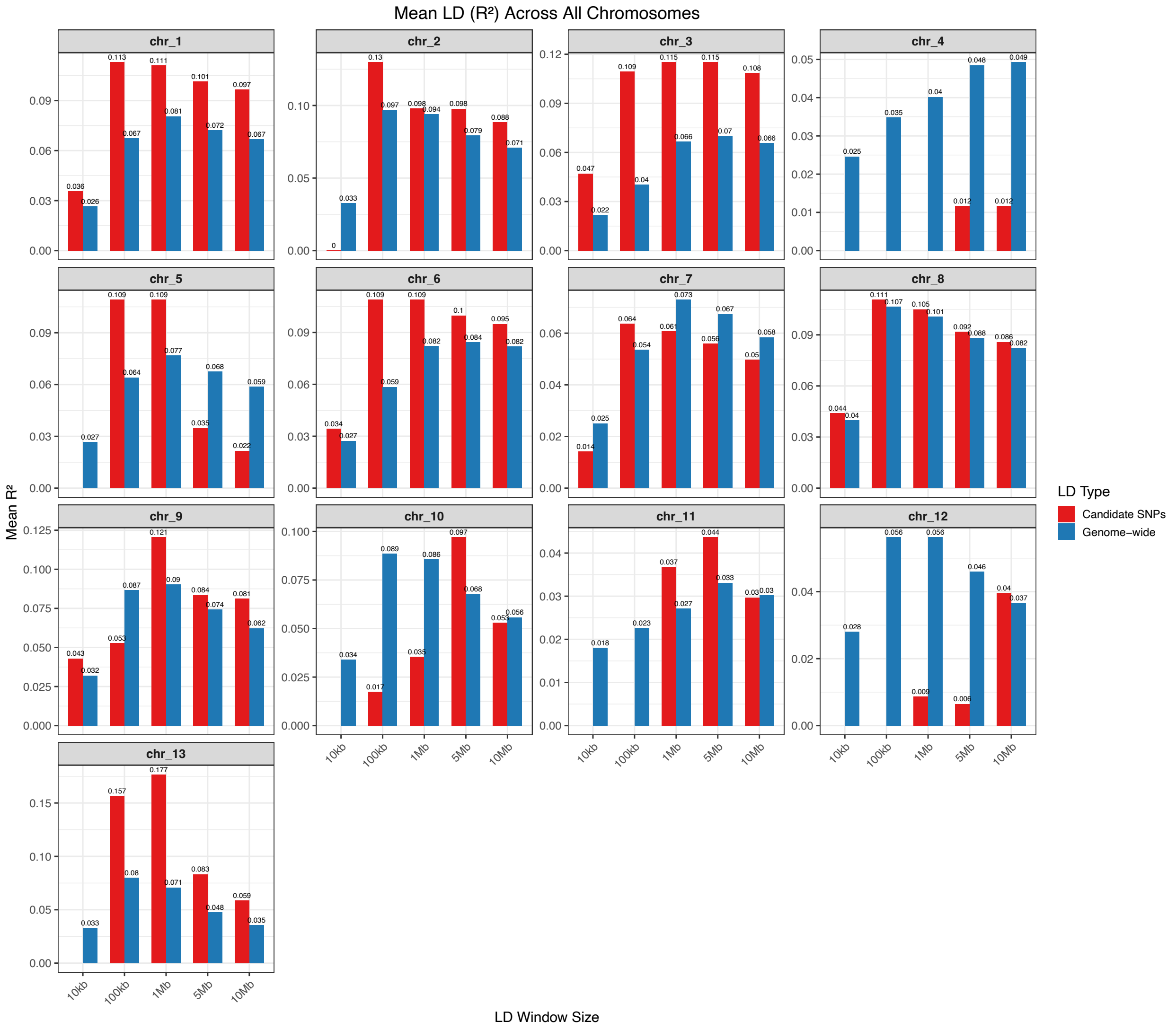


Fig. S9. Genome-wide comparison of linkage disequilibrium (LD) between candidate and background SNPs across chromosomes and distance windows for Shanghai individuals. Barplots show the mean pairwise LD ($R^2$) for SNPs identified as outliers by PCAdapt ("Candidate SNPs") versus all genome-wide SNPs ("Genome-wide") across five genomic distance windows (10 kb, 100 kb, 1 Mb, 5 Mb, and 10 Mb), computed separately for each chromosome. LD was calculated using VCFtools for our final dataset with 158,235 SNPs, and 7318 candidate SNPs identified by PCAdapt. On several chromosomes candidate SNPs exhibit substantially higher LD than the genome-wide background, particularly at intermediate window sizes (100 kb to 1 Mb). Chromosomes 1, 2, 3, 5, 6, 9, and 13 stood out as having candidate LD values that exceeded background levels.

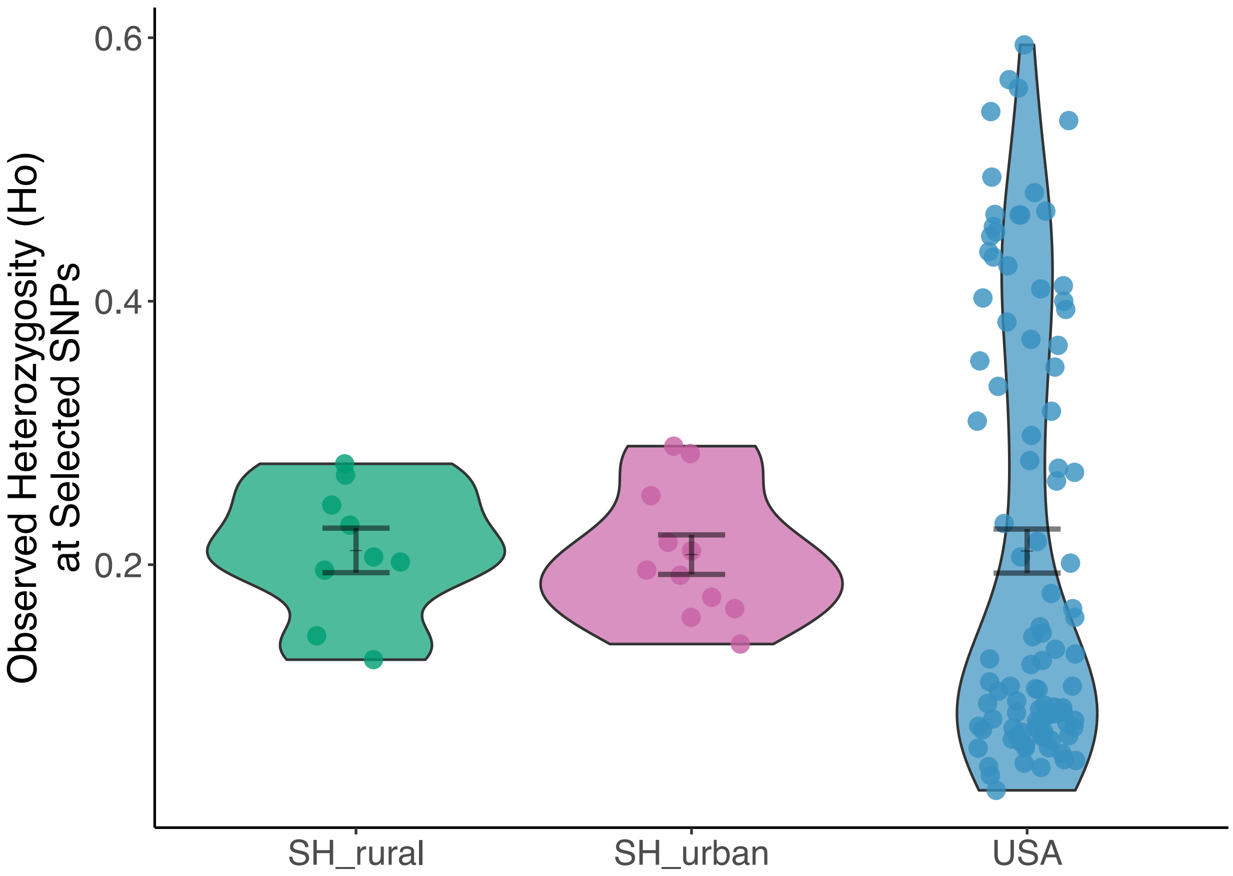

Fig. S10. Observed heterozygosity (Ho) at 184 candidate SNPs is similar across native and invasive populations. Violin show observed heterozygosity at 184 candidate SNPs identified as under selection in Shanghai rural (n = 9), Shanghai urban (n = 11), and invasive USA populations (n = 98). Heterozygosity at putatively adaptive loci is comparable across all groups (USA: 0.210 ± 0.017; SH rural: 0.211 ± 0.017; SH urban: 0.208 ± 0.015).

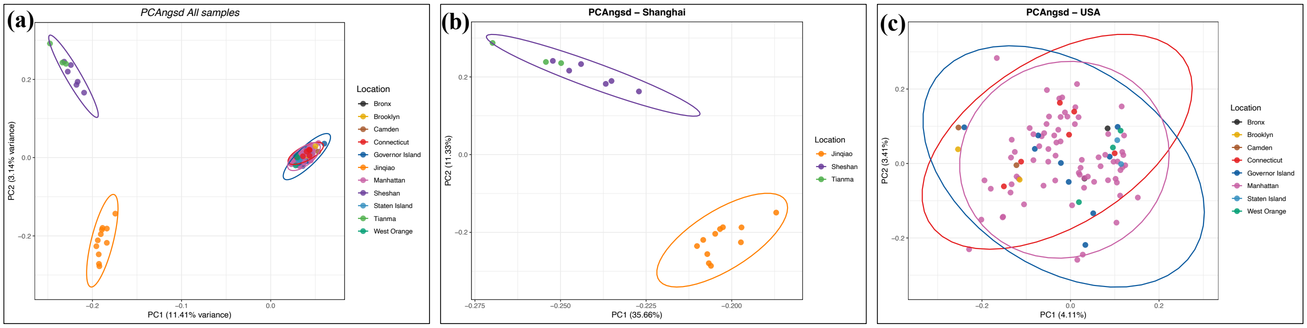


Fig. S11. Principal component analyses using PCAngsd across different locations. (a) PCA of all samples showing clear separation between native Chinese (Jinqiao, Sheshan, Tianma) and invasive USA populations along PC1 (11.41% variance), and Chinese samples separating along PC2 by habitat type (3.14% variance). (b) PCA of Shanghai individuals (PC1: 35.66%, PC2: 11.33% variance), with clear separation between Jinqiao (orange), Sheshan (purple), and Tianma (green) locations. (c) PCA of USA individuals revealing minimal genetic differentiation across locations (PC1: 4.11%, PC2: 3.41% variance). Colored ellipses represent 95% confidence intervals for each population group.

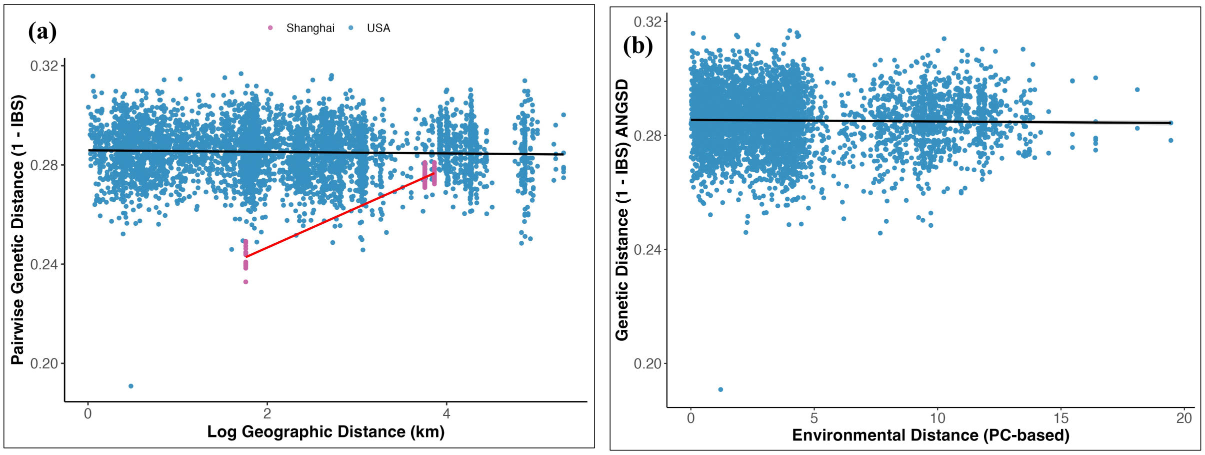


Fig. S12. Isolation-by-distance (IBD) and isolation-by-environment (IBE) analyses based on pairwise genetic distances calculated using genotype likelihoods in ANGSD. (a) Relationship between pairwise genetic distance (1 – IBS) and log-transformed geographic distance (km) across urban and rural populations in the United States (blue) and Shanghai, China (pink). Solid black and red lines indicate fitted linear regressions for USA and Shanghai, respectively. (b) Relationship between pairwise genetic distance and environmental distance (PC-based) across USA populations. Despite differences in absolute genetic distance values, the overall trends were consistent with the PLINK-based analyses.

Table S1: Demographic modelling parameter ranges and demographic events modeled across eight alternative scenarios, including estimates of population sizes, founder bottlenecks, and timing of population splits used in final dataset with

158,235 SNPsMomi2. See Supplementary figure S7 for model details.

| Parameter | Model 1 | Model 2 | Model 3 | Model 4 | Model 5 | Model 6 | Model 7 | Model 8 |
|---|---|---|---|---|---|---|---|---|
| Initial N_e | 1.90E+06 | 1.90E+06 | 1.90E+06 | 1.90E+06 | 1.90E+06 | 1.90E+06 | 1.90E+06 | 1.90E+06 |
| N_rural | (1e3, 1e7) | (1e3, 1e7) | (1e3, 1e7) | (1e3, 1e7) | (1e3, 1e7) | (1e3, 1e7) | (1e3, 1e7) | (1e3, 1e7) |
| N_urban | (1e3, 1e7) | (1e3, 1e7) | (1e3, 1e7) | (1e3, 1e7) | (1e3, 1e7) | (1e3, 1e7) | (1e3, 1e7) | (1e3, 1e7) |
| N_usa | (1e3, 1e7) | (1e3, 1e7) | (1e3, 1e7) | (1e3, 1e7) | (1e3, 1e7) | (1e3, 1e7) | (1e3, 1e7) | (1e3, 1e7) |
| N_ancestral | N/A | N/A | N/A | N/A | (1e3, 1e7) | (1e3, 1e7) | (1e3, 1e7) | (1e3, 1e7) |
| N_urban_founder | (1e1, 1e3) | (1e1, 1e3) | (1e1, 1e3) | (1e1, 1e3) | N/A | N/A | N/A | N/A |
| N_korea_founder | (1e1, 1e3) | N/A | (1e1, 1e3) | N/A | (1e1, 1e3) | N/A | (1e1, 1e3) | N/A |
| N_usa_founder | (1e1, 1e3) | 1e1, 1e3) | (1e1, 1e3) | 1e1, 1e3) | (1e1, 1e3) | 1e1, 1e3) | (1e1, 1e3) | (1e1, 1e3) |
| t_usa_split | (8, 20), t0=10 | (8, 20), t0=10 | (8, 20), t0=10 | (8, 20), t0=10 | (8, 20), t0=10 | (8, 20), t0=10 | (8, 20), t0=10 | (8, 20), t0=10 |
| t_korea_split | (t_usa_split, 30), t0=16 | N/A | (t_usa_split, 30), t0=16 | N/A | (t_usa_split, 30), t0=16 | N/A | (t_usa_split, 30), t0=16 | N/A |
| t_urban_split | (t_korea_split, 1e6), t0=50 | (t_usa_split, 1e6), t0=50 | (t_korea_split, 1e6), t0=50 | (t_usa_split, 1e6), t0=50 | N/A | N/A | N/A | N/A |
| t_rural_urban_split | N/A | N/A | N/A | N/A | (t_korea_split, 1e6), t0=50 | (t_usa_split, 1e6), t0=50 | (t_korea_split, 1e6), t0=50 | (t_usa_split, 1e6), t0=50 |
| Invasion Source to USA | Urban via Korea | Urban directly | Rural via Korea | Rural directly | Urban via Korea | Urban directly | Rural via Korea | Rural directly |

Table S2: AIC-based model comparison for demographic inference of the Spotted Lanternfly invasion. Summary statistics for eight alternative demographic models evaluated using Akaike Information Criterion (AIC) across 785 bootstrap replicates. The best-supported model is Model5_Independent_Urban_Korea, with the lowest mean AIC (2,983,862.169), smallest ΔAIC (5,233.553), and the highest mean AIC weight (0.936). Confidence intervals (CI95) are shown for mean AIC scores to indicate the uncertainty across bootstrap replicates.

| Model | Mean_AIC | SD_AIC | Mean_LogLikelihood | Mean_DeltaAIC | Mean_AICWeight | N | CI95_AIC_Lower | CI95_AIC_Upper |
|---|---|---|---|---|---|---|---|---|
| **Model5_Independent_Urban_Korea** | **2983862.169** | **41697.991** | **-1491922.085** | **5233.553** | **0.936** | **785.000** | **2980940.713** | **2986783.626** |
| Model1_Urban_Korea | 3005380.315 | 42677.790 | -1502681.157 | 26751.699 | 0.054 | 785.000 | 3002390.212 | 3008370.418 |
| Model7_Independent_Rural_Korea | 3008176.349 | 53196.899 | -1504079.175 | 29547.733 | 0.010 | 785.000 | 3004449.253 | 3011903.445 |
| Model3_Rural_Korea | 3018696.409 | 57363.756 | -1509339.205 | 40067.793 | 0.000 | 785.000 | 3014677.374 | 3022715.444 |
| Model6_Independent_Urban_Direct | 3555869.781 | 40550.015 | -1777927.891 | 577241.165 | 0.000 | 785.000 | 3553028.755 | 3558710.807 |
| Model2_Urban_Direct | 3593089.548 | 52607.171 | -1796537.774 | 614460.932 | 0.000 | 785.000 | 3589403.770 | 3596775.326 |
| Model8_Independent_Rural_Direct | 3627913.549 | 76231.105 | -1813949.775 | 649284.933 | 0.000 | 785.000 | 3622572.625 | 3633254.474 |
| Model4_Rural_Direct | 3730117.939 | 79771.462 | -1865051.969 | 751489.322 | 0.000 | 785.000 | 3724528.968 | 3735706.909 |

Table S3: Statistics for all parameters for the best model (Model5_Independent_Urban_Korea) with bootstrap analysis (735 independent replicates) providing confidence intervals for model parameters.

| | mean | sd | min | max | lower_CI | upper_CI |
|---|---|---|---|---|---|---|
| t_usa_split | 8.952 | 2.970 | 8.000 | 20.000 | 8.737 | 9.167 |
| t_korea_split | 29.686 | 0.862 | 19.650 | 30.000 | 29.624 | 29.748 |
| t_rural_urban_split | 171.427 | 7.042 | 149.945 | 226.034 | 170.917 | 171.937 |
| N_rural | 1001.682 | 30.414 | 1000.000 | 1750.379 | 999.479 | 1003.884 |
| N_urban | 1013.231 | 123.007 | 1000.000 | 2780.220 | 1004.323 | 1022.138 |
| N_usa | 1953.236 | 4497.003 | 1000.000 | 63576.261 | 1627.591 | 2278.881 |
| N_ancestral | 2828.105 | 82.068 | 2201.860 | 3830.685 | 2822.162 | 2834.048 |
| N_korea_founder | 46.028 | 7.525 | 14.717 | 51.562 | 45.483 | 46.573 |
| N_usa_founder | 514.496 | 299.154 | 10.846 | 997.846 | 492.834 | 536.159 |

Table S4: Estimated time to most recent common ancestor (TMRCA) calculated from ROHs in native (CN) and invasive (USA) populations of Spotted Lanternfly based on median ROH lengths within three size classes and assuming a recombination rate of 1 cM/Mb (1 generation/year).

| Country | Size_class | Median_length | Recombination Rate | TMRCA_years |
|---|---|---|---|---|
| CN | 0.5-1 Mb | 0.625 | 1 cM/Mb | 80 |
| CN | 1-5 Mb | 1.17 | 1 cM/Mb | 42.7 |
| USA | 0.5-1 Mb | 0.660 | 1 cM/Mb | 75.8 |
| USA | 1-5 Mb | 1.38 | 1 cM/Mb | 36.2 |
| USA | >5 Mb | 5.62 | 1 cM/Mb | 8.9 |

Table S5. SNP density across chromosomes in Spotted Lanternflies of the USA population from our final VCF. SNP density is reported as the number of SNPs per megabase (SNPs/Mb).

| Chromosome | SNPs | Chr_Length (kb) | SNPs/Mb |
|---|---|---|---|
| chr_1 | 30,292 | 404,012 | 75 |
| chr_2 | 16,974 | 238,731 | 71.1 |
| chr_3 | 19,418 | 227,283 | 85.4 |
| chr_4 | 3,575 | 217,703 | 16.4 |
| chr_5 | 13,881 | 182,427 | 76.1 |
| chr_6 | 11,852 | 165,984 | 71.4 |
| chr_7 | 13,131 | 149,133 | 88.1 |
| chr_8 | 12,382 | 142,606 | 86.9 |
| chr_9 | 8,976 | 133,192 | 67.4 |
| chr_10 | 9,769 | 123,181 | 79.3 |
| chr_11 | 7,415 | 83,141 | 89.2 |
| chr_12 | 6,464 | 78,192 | 82.7 |
| chr_13 | 4,106 | 55,467 | 74 |
